# Ancestral hybridisation facilitated species diversification in the Lake Malawi cichlid fish adaptive radiation

**DOI:** 10.1101/738633

**Authors:** Hannes Svardal, Fu Xiang Quah, Milan Malinsky, Benjamin P Ngatunga, Eric A Miska, Walter Salzburger, Martin J Genner, George F Turner, Richard Durbin

## Abstract

The adaptive radiation of cichlid fishes in East Afrian Lake Malawi encompasses over 500 species that are believed to have evolved within the last 800 thousand years from a common founder population. It has been proposed that hybridisation between ancestral lineages can provide the genetic raw material to fuel such exceptionally high diversification rates, and evidence for this has recently been presented for the Lake Victoria Region cichlid superflock. Here we report that Lake Malawi cichlid genomes also show evidence of hybridisation between two lineages that split 3-4 million years ago, today represented by Lake Victoria cichlids and the riverine *Astatotilapia* sp. ‘ruaha blue’. The two ancestries in Malawi cichlid genomes are present in large blocks of several kilobases, but there is little variation in this pattern between Malawi cichlid species, suggesting that the large-scale mosaic structure of the genomes was largely established prior to the radiation. Nevertheless, tens of thousands of polymorphic variants apparently derived from the hybridisation are interspersed in the genomes. These loci show a striking excess of differentiation across ecological subgroups in the Lake Malawi cichlid assemblage, and parental alleles sort differentially into benthic and pelagic Malawi cichlid lineages, consistent with strong differential selection on these loci during species divergence. Furthermore, these loci are enriched for genes involved in immune response and vision, including opsin genes previously identified as important for speciation. Our results reinforce the role of ancestral hybridisation in explosive diversification by demonstrating its significance in one of the largest recent vertebrate adaptive radiations.

## Introduction

Adaptive radiation, the rapid diversification of one or a few ancestral lineages into a number of species that occupy a range of ecological niches, is thought to be responsible for a considerable fraction of extant biodiversity (Berner and Salzburger 2015). A growing body of research investigates the intrinsic and extrinsic factors that may facilitate adaptive radiation (reviewed in Seehausen 2015). This may be addressed by correlating ecological factors and organismal features with the propensity for adaptive diversification across taxa (Wagner et al. 2012), but the ultimate goal must be to gain a functional understanding of the genetic and ecological mechanisms at play.

The role of hybridisation in adaptive radiation and speciation has been debated (Mallet 2007; Abbott et al. 2013; Schumer et al. 2014). While the cessation of genetic exchange between sister lineages is often seen as a prerequisite for speciation, it has been proposed that both hybridisation into the ancestral lineage as well as genetic exchange between diverging lineages within a radiation could favour adaptive diversification (Seehausen 2004; Seehausen 2013; Marques et al. 2019). With genome-wide sequencing data becoming increasingly accessible, the latter scenario of genetic exchange between diverging lineages has been shown to be widespread in plants and animals (Mallet et al. 2015; Novikova et al. 2016; Svardal et al. 2017), especially in young adaptive radiations (Dowling and DeMarais 1993; Heliconius Genome Consortium 2012; Lamichhaney et al. 2015; Stryjewski and Sorenson 2017; Edelman et al. 2018; Kozak et al. 2018; Malinsky et al. 2018), and links to adaptation have been made (Heliconius Genome Consortium 2012; Richards and Martin 2017; Malinsky et al. 2018). However, the generality and evolutionary importance of hybridisation in the seeding population of an adaptive radiation, that is, the ancestral lineage before initial divergence, is less well understood.

Hybridisation in the common ancestor of an adaptive radiation could promote adaptive radiation in several ways (reviewed in Seehausen 2004; Marques et al. 2019). Hybrids often exhibit novel or extreme characters, known as transgressive segregation (Rieseberg et al. 1999). In particular, epistatic interactions of alleles or – at a higher level – genetic modules from two or more parental backgrounds can lead to novel phenotypes (Dittrich-Reed and Fitzpatrick 2013) that could confer a fitness advantage in previously underutilized ecological niches or provide a target for sexual selection. However, even in the absence of interactions between loci, hybridisation is expected to increase the genetic variance of a quantitative trait, if alleles of different effect sizes and directions segregate in the two parental populations. This is true even if the parental lineages were (independently) under stabilising selection for the same trait optimum. The heritable genetic variability produced in this way could facilitate adaptive divergence. Furthermore, genetic incompatibility, that is, negative epistasis between loci fixed for different alleles in the parental lineages, could potentially provide an axis of divergent selection (Schumer et al. 2015) that couples with and reinforces natural divergent selection, but whether this actually happens in nature and – if it does – how common it is, is largely an open question.

Recent advances in genome sequencing have provided empirical evidence that ancestral hybridisation can occur prior to adaptive radiation (Barrier et al. 1999) and can generate phenotypic novelty (Stryjewski and Sorenson 2017). A group of species that has received considerable attention with respect to their repeated rapid adaptive diversification and the role of hybridisation in this process are haplochromine cichlids, a tribe of percomorph fish widely-distributed in eastern Africa including the species of the adaptive radiations in the Lake Victoria region, in Lake Malawi, and a subset of the species of the adaptive radiation of cichlid fishes in Lake Tanganyika (Kocher 2004; Salzburger 2018). Meier et al. (2017) demonstrated that the haplochromine cichlid fishes collectively known as the Lake Victoria Region Superflock (LVRS) emerged from a hybrid ancestor, that loci highly divergent across LVRS lineages are enriched for alleles differentiated between the two ancestral lineages, and that ecologically divergent opsin alleles originate from the two ancestors. Furthermore, hybridisation was also reported between ancestral lineages of Lake Tanganyika cichlids (Irisarri et al. 2018) as well as between some of the main lineages (‘tribes’) in this lake (Meyer et al. 2016). Ancient hybridisation has also been suggested to play a role in the Lake Malawi cichlid adaptive radiation (Joyce et al. 2011; Genner and Turner 2012), but the suggested events were based on mitochondrial evidence, which was not supported by recent whole-genome sequencing analyses (Malinsky et al. 2018).

Here we re-investigate the occurrence and role of ancestral hybridisation in the Lake Malawi cichlid adaptive radiation by analysing recently published and newly obtained whole-genome sequences of haplochromine cichlids from Lake Malawi, the LVRS, as well as other East African lakes and rivers (Fig. 1; supplementary table 1) (Brawand et al. 2014; McGee et al. 2016; Malinsky et al. 2018). We find strong evidence for ancient hybridisation at the base of the Malawi cichlid radiation, between a lineage related to the LVRS and a riverine lineage previously shown to carry a Malawi-like mitochondrial haplotype (Genner et al. 2015). Furthermore, we show that the majority of the genetic variation produced by this hybridisation event has been lost in Malawi by the relatively rapid fixation at most loci of one or the other parental contribution in the ancestral population of Malawi cichlids. The genetic variation created by this hybridisation event that still persists in Malawi shows elevated divergence across the early splits in the radiation, beyond its effect of increasing minor allele frequencies, and is enriched in gene ontologies related to vision and the immune system.

**FIG. 1.**
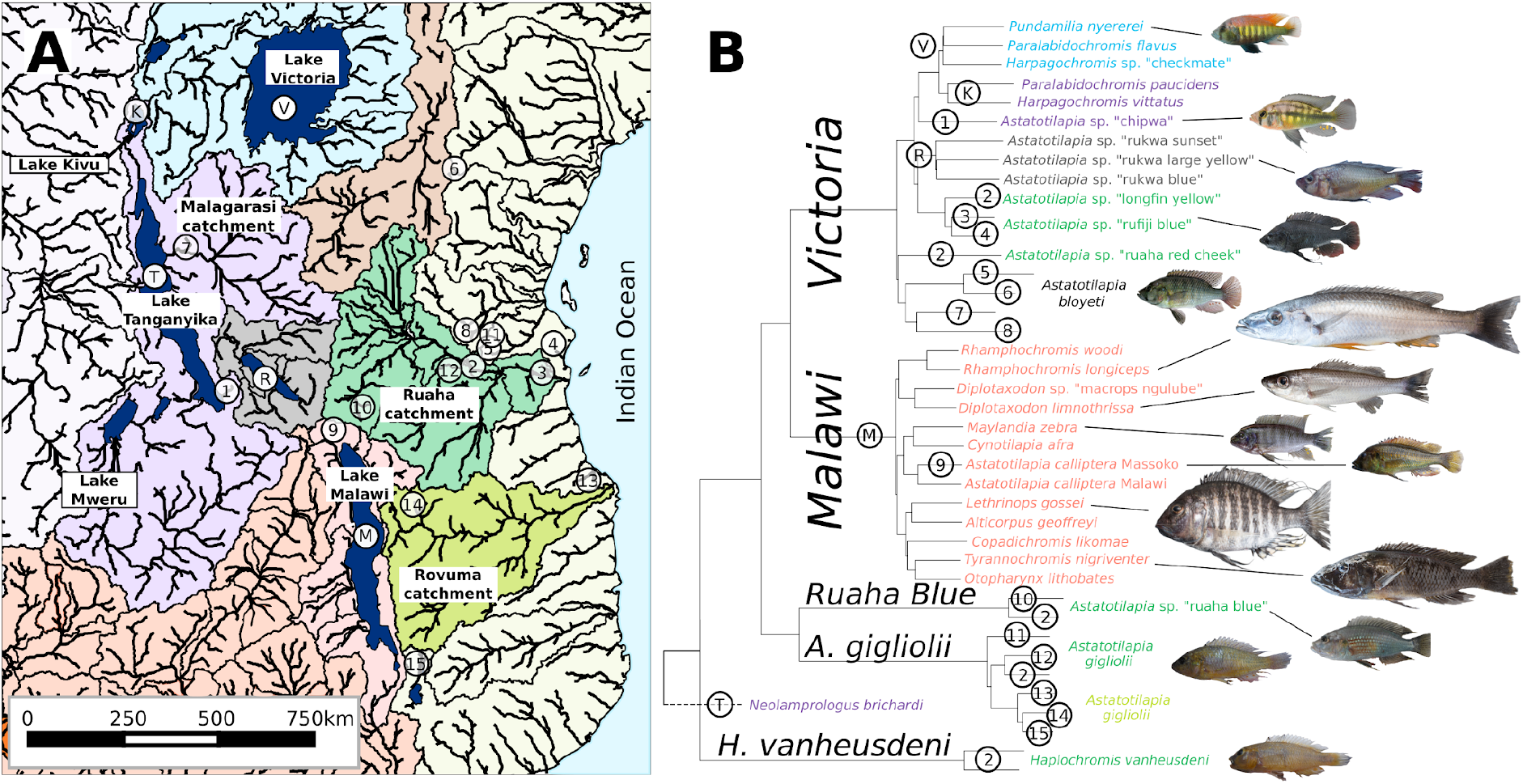
Sampling and genetic relationships. (a) Map of East African river catchments and sampling locations. (b) Neighbour-joining tree of pairwise genetic distances between all samples. Colours correspond to river catchments in (a). Black italicised clade names are used in the text to refer to the clades. The *Victoria* clade contains the Lake Victoria Region Superflock (LVRS) and widely-distributed riverine haplochromines. Some members of the *A. gigliolii* clade were previously referred to as *A. tweddlei* but after examining type specimens we suggest this to be a junior synonym of *A. gigliolii*. Relative scaling of fish pictures is to their approximate body sizes. Image credit *P. nyererei*: O. Selz. Morte details on samples, sampling locations and fish images can be found in supplementary table 1.

## Results

### Sampling, sequencing, and variant detection

We obtained samples for representatives of the haplochromine cichlid adaptive radiations in the Lake Victoria region, from Lake Malawi, and from riverine and lacustrine haplochromines that have been proposed to be close relatives of these radiations (fig. 1a, supplementary table 1, Materials and Methods) (Genner et al. 2015; Malinsky et al. 2018). The Lake Malawi sampling comprised two samples for each of the previously defined major eco-morphological groups (only one sample for *utaka*, supplementary table 1) (Malinsky et al. 2018). Illumina short read sequence data was partly obtained from previous studies (Brawand et al. 2014; McGee et al. 2016; Malinsky et al. 2018) and partly sequenced for this study (supplementary table 1). DNA libraries were created and whole-genome sequenced on an Illumina HiSeq platform to individual coverages between 7.9 and 19.2 fold (median 14.3 fold). Reads were aligned to the Nile tilapia reference genome Orenil. 1.1 (GCA_000188235.2) (Brawand et al. 2014) and variants called. After filtering, 19 million biallelic SNP variants were kept for further analysis (Materials and Methods).

### Gene flow into the ancestral population of the Lake Malawi cichlid flock

To provide an overview of genetic relationships between all samples we first built a neighbour-joining (NJ) tree from pairwise genetic distances (fig. 1b). The tree suggests that the sister lineage to the Lake Malawi species flock (*Malawi* in the following) is a large clade containing widely distributed riverine haplochromines as well as the Lake Victoria Region Superflock (collectively called *Victoria*, in the following).

Based on mitochondrial DNA evidence, Genner et al. (2015) previously suggested that *Astatotilapia* sp. ‘ruaha blue’ (*Ruaha Blue* in the following) from the Ruaha catchment constitutes a sister lineage to the Lake Malawi radiation. Although whole-genome data does not support this suggestion of a sister species relationship between the taxa (fig. 1b), calculation of Patterson’s D (ABBA-BABA test) (Patterson et al. 2012) revealed a substantial excess of allele sharing between Lake Malawi samples and *Ruaha Blue* relative to *Victoria* (fig. 2a). Using a combination of D tests similar to the 5-taxon test proposed by Pease and Hahn (2014), we further confirmed that this excess allele sharing is most likely due to introgression from the *Ruaha Blue* lineage into the *Malawi* lineage (fig. 2b). In particular, the strongest signals correspond to excess allele sharing between *Ruaha Blue* and *Malawi* relative to *Victoria* (fig. 2b, red) and the second strongest to excess allele sharing between *Malawi* and *Ruaha Blue* relative to *A. gigliolii* (fig. 2b, green). This suggests that the lineages leading to present day *Ruaha Blue* and *Malawi* exchanged genes. Furthermore, we observe a significant excess of allele sharing between *A. gigliolii* and *Malawi* relative to *Victoria* (fig. 2b, turquoise), which is expected under gene flow from *Ruaha Blue* into *Malawi* but not for the opposite direction. Finally, the weakest signal is an excess of allele sharing between *Ruaha Blue* and *Victoria* relative to *A. gigliolii* (fig. 2b, blue). This signal is significant for most comparisons of samples within the clades but not for all. While this signal cannot be explained by gene flow from *Ruaha Blue* into *Malawi*, we attribute this to further genetic exchange, for example between riverine haplochromines in the *Victoria* clade and *Ruaha Blue*, because the signal shows clear geographic structure, with the strongest excess of allele sharing between the sympatric *Ruaha Blue* and *A.* sp. ‘ruaha red cheek’ from the *Victoria* clade.

**FIG. 2.**
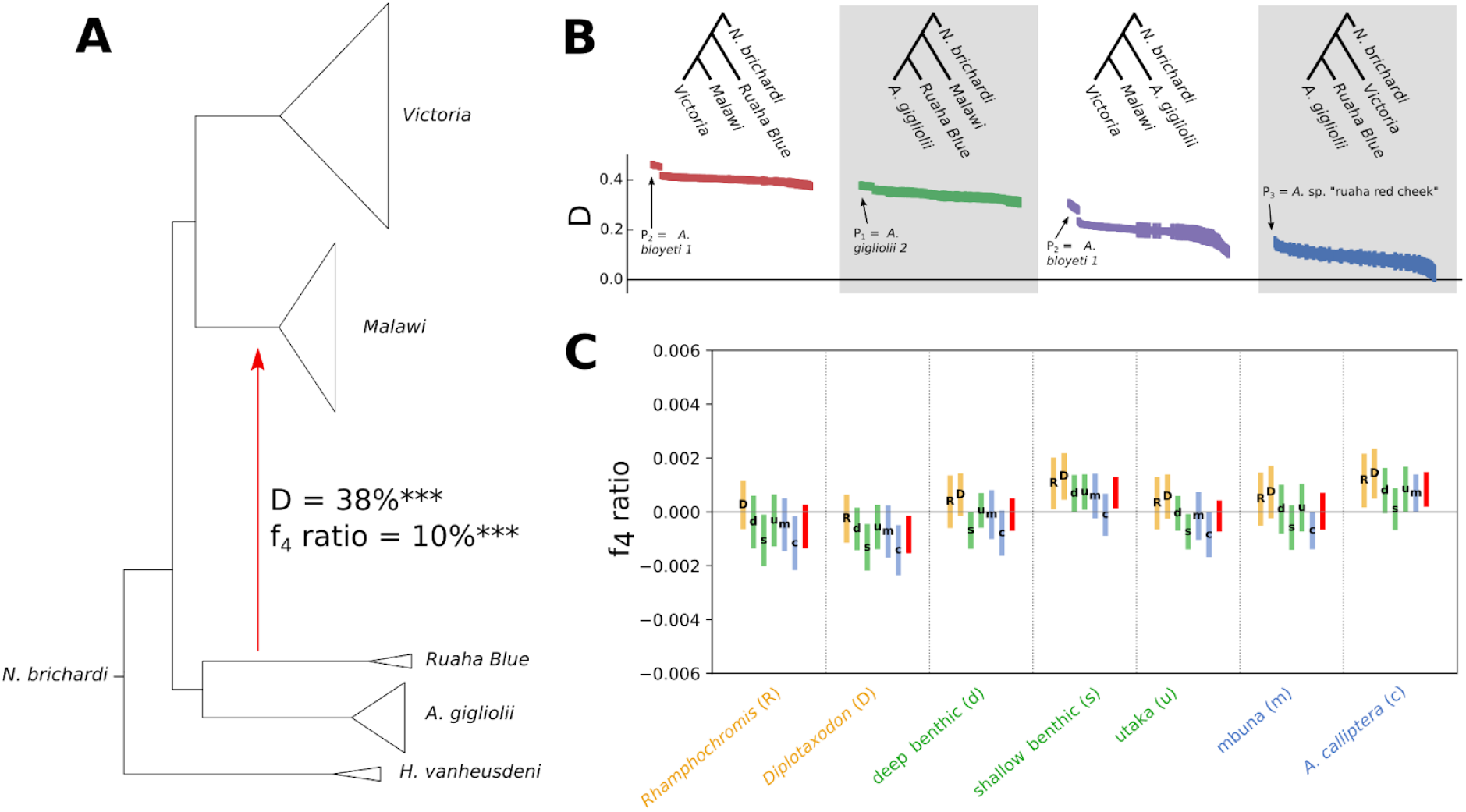
Gene flow into the common ancestor of the Malawi radiation (a) Cladogram of major lineages with the red arrow indicating the strongest gene flow event as inferred by Patterson’s D and f_4_ admixture ratio (P_1_ = *Victoria*, P_2_ = *Malawi*; P_3_ = *Ruaha Blue*, P_4_ = *N. brichardi*) (Patterson et al. 2012). ***indicates a block-jackknifing significance of p<10^−16^. (b) D tests to infer the directionality of introgression. The four regions accentuated by white and grey shading correspond to -D_FI_, D_IL_, -D_OL_ and D_FO_, respectively, in the D_FOIL_ 5-taxon test (*Victoria*, *Malawi*, *Ruaha Blue*, *A. gigliolii*, *N. brichardi)* (Pease and Hahn 2014), except that – as is standard for Patterson’s D – only sites were considered where P_3_ has the derived allele. The trees within each region show the groups from which P_1_, P_2_, P_3_, and P_4_ (from left to right) were taken for the tests below. The different data points correspond to tests for all possible combinations of samples from each of the clades. Error bars correspond to ± three block-jackknifing standard deviations. Outlier tests are annotated with the sample they have in common. We note that the sample *A. bloyeti 1* from Lake Kumba in the Pangani catchment is an outlier in many comparisons consistent with it having received genetic material not related to any other sample in our dataset. (c) f_4_ admixture ratio tests (Patterson et al. 2012) with P_3_ = *Victoria* and P_4_ = *Ruaha Blue*. P_2_ is given on the x-axis and P_1_ by the letter in the bar. Bars and labels are colored by group microhabitat (yellow = pelagic, green = benthic, blue = littoral) The red bars correspond to the case where P_1_ is a group of all *Malawi* samples except P_2_. Bar height corresponds to ± three block-jackknifing standard deviations. f_4_ values between Malawi subgroups in (c) are a hundred times smaller than between founders in (a).

Using the f_4_ admixture ratio (f_4_ ratio) (Patterson et al. 2012), we estimated that on average ∼10% of *Malawi* genetic material traces its ancestry through the gene flow event from *Ruaha Blue* (fig. 2a). However, we note that in coalescent simulations this estimate is biased downwards: an f_4_ admixture ratio estimate of 10% was compatible with a model with 22% actual gene flow (supplementary figs. S1, S2). Using a coalescent-based approach, we estimate the split time between *Victoria* and *Ruaha Blue* lineages to 3.2-3.9 million years ago and the split of *Victoria* and *Malawi* to 2.0-2.8 million years ago (supplementary fig. S3).

The Lake Malawi cichlid assemblage can be organised into six genetically well-defined groups (Malinsky et al. 2018). To test whether different *Malawi* groups share different fractions of *Ruaha Blue* alleles, we calculated f_4_ ratios of the form f_4_(P_1_=*Malawi*-1, P_2_=*Malawi*-2; P_3_=*Victoria*, P_4_=*Ruaha Blue*), where *Malawi*-1 and *Malawi*-2 are pairs of *Malawi* groups (fig. 2c). Such tests are expected to be significantly positive if *Malawi*-1 shares significantly more ancestry with *Ruaha Blue* than *Malawi*-2 (or reciprocally negative if it shares less ancestry). These tests were mostly not significant and even those that are marginally significant are of low magnitude <0.1%, a hundred times lower than f_4_(P_1_=*Victoria*, P_2_=*Malawi*; P_3_=*Ruaha Blue*, P_4_=*Outgroup*). From this, we conclude that gene flow most likely occurred in the ancestral population prior to the radiation of the present-day *Malawi* groups and that overall there is no strong differential retention of *Victoria*/*Ruaha Blue*-specific alleles across *Malawi* lineages. *Victoria* and *Ruaha Blue* thus constitute extant relatives of two ancestral lineages that contributed to the Lake Malawi radiation.

### Introgressed polymorphisms show excess divergence across the Malawi radiation

To test whether alleles that introgressed from the *Ruaha-Blue*-related lineage into the *Malawi* ancestor played a role in species divergence, we examined whether introgression-derived variants are enriched for high divergence across Malawi eco-morphological groups, adopting an approach similar to that used by Meier et al. (2017). We split variants in our data set that are polymorphic within *Malawi* into four categories depending on their allelic states and variation patterns in *Victoria*, *Ruaha Blue* and an outgroup. The categories comprised (1) variants that are fixed for the same allele in *Victoria* and *Ruaha Blue*, (2) variants for which both *Malawi* alleles are also seen in *Victoria and/or Ruaha Blue*, (3) variants with the alternative alleles differentially fixed between *Victoria* and *Ruaha Blue*, and (4) variants that are fixed for the same allele in *Victoria* and *Ruaha Blue* but where the alternative allele is fixed in the outgroup *‘H’. vanheusdeni*, respectively (table 1).

**Table 1.**
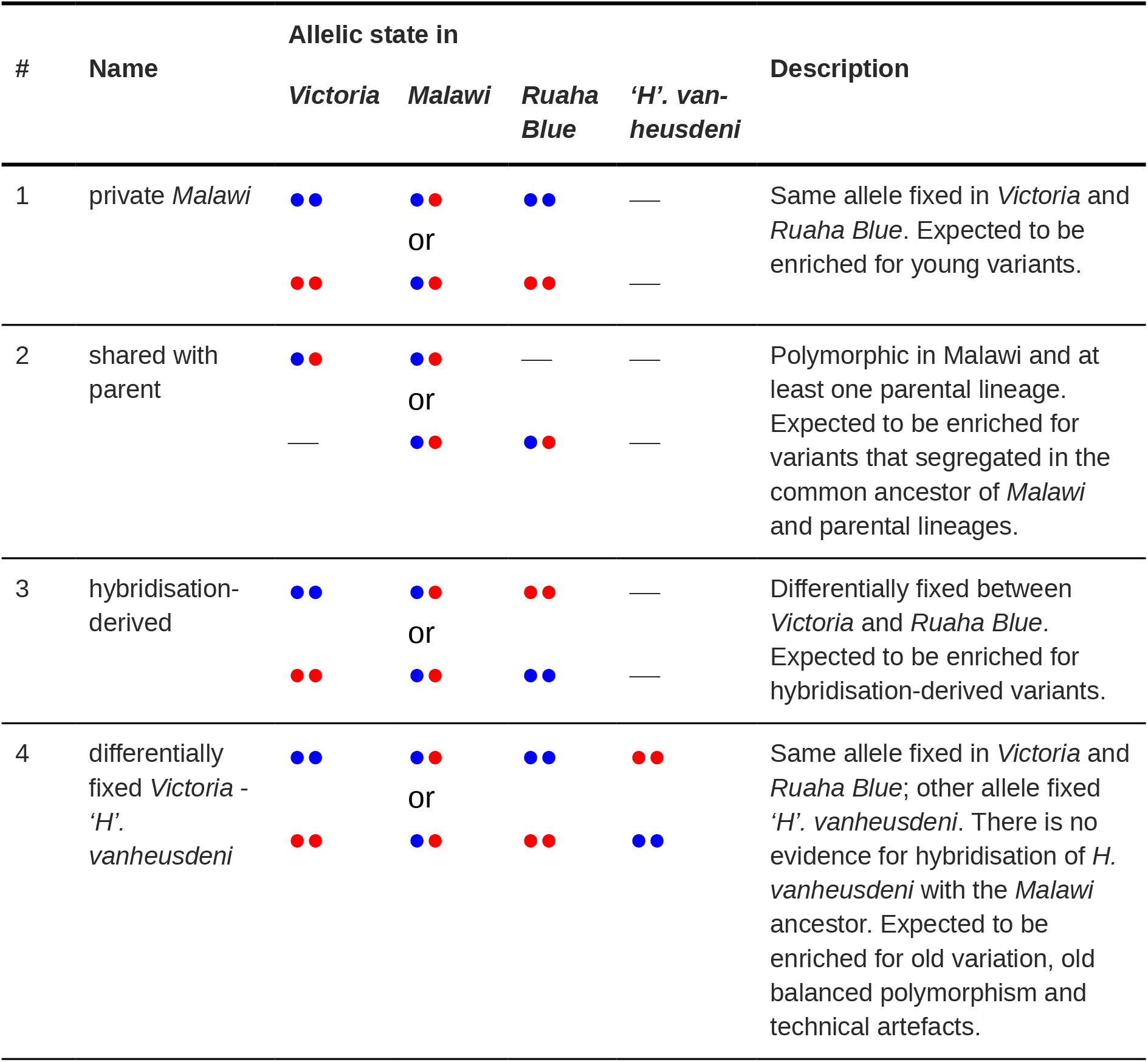
Description of the four variant categories used for the analyses shown in fig. 3.

Each of these groups is expected to be enriched for different types of variants. The first and by far the largest group is enriched for young variants that arose within *Malawi* or on the ancestral branch leading to *Malawi*, while the second group represents shared variation that is enriched for old variants that segregated in the common ancestor of *Malawi* and *Victoria*. The third group is enriched for variants for which the polymorphism in *Malawi* was created by alleles that diverged on the branches separating the two ancestral lineages. We will therefore refer to variants in this category as "hybridisation-derived". Since the third pattern could in principle also be created by technical artefacts or by old polymorphism that was segregating all the way along the branch from the common ancestor of *Malawi*, *Victoria*, and *Ruaha Blue* down to *Malawi*, we added the fourth "control" category, which is expected to include both artefacts and very old polymorphisms. *‘H’. vanheusdeni* used for this control category does not show any signs for introgression into the *Malawi* ancestor; on the contrary, it shows significant excess allele sharing with Victoria compared to Malawi, albeit with a signal an order of magnitude smaller than the one involving *Ruaha Blue* reported above (D(*Malawi*, *Victoria*; *‘H’. vanheusdeni*, *N. brichardi*) = 0.04).

Next, we identified SNPs that are highly differentiated between the pelagic (*Rhamphochromis*, *Diplotaxodon*) and benthic (all other *Malawi*) lineages (*pelagic* and *benthic*, in the following) within the Lake Malawi cichlid adaptive radiation, which was suggested to be first split in the radiation (Malinsky et al. 2018) (see also fig. 1), by taking the top 1% of SNPs ranked by fixation index (F_ST_) value, a relative measure of genetic divergence. For each of the above mentioned categories we then asked what proportion of SNPs were highly differentiated (fig. 3b). We found that the set of hybridisation-derived variants (category 3) is more than three times more likely to be among F_ST_ outliers compared with the set of all variants and 2.4 times more likely compared with variants shared with a parent, which represents a significant enrichment (fig. 3b; Welch’s t-test p=10^−13^ and 10^−11^, respectively). While variants shared with a parental lineage (category 2) and variants differentially fixed between *Victoria* and *‘H’. vanheusdeni* (category 4) also showed an excess of divergent loci compared to private *Malawi* variants, the effect was much less pronounced (1.44 and 1.16 times for categories 2 and 4 respectively, compared to 3.5 times for category 3). Similar results were obtained with different choices of F_ST_ outlier thresholds (supplementary fig. S4), and also if the variant categories were ascertained with equally sized samples from *Victoria* and *Ruaha Blue* (supplementary fig. S5).

**FIG. 3.**
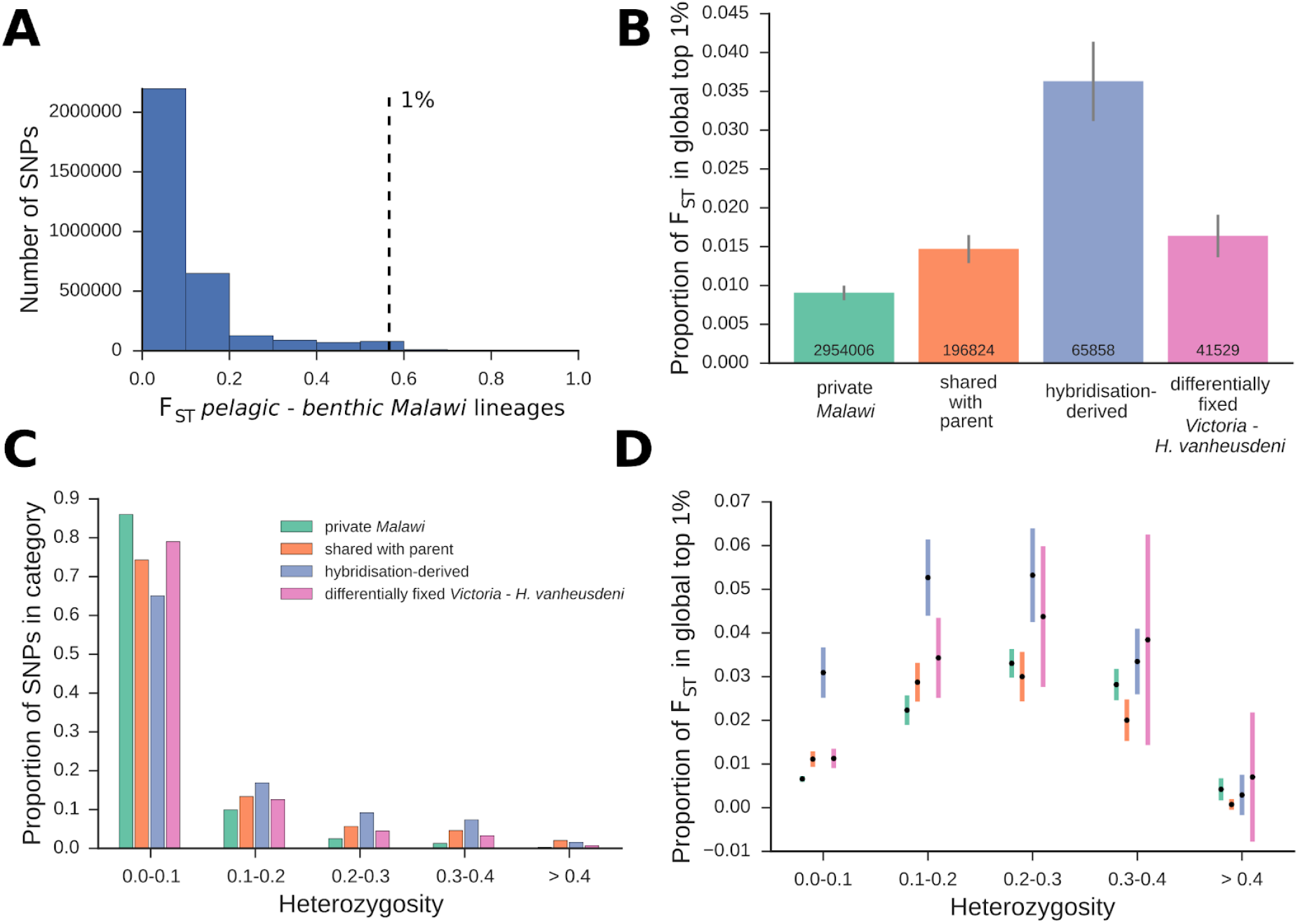
Distribution of F_ST_ outliers across variant categories described in table 1. (a) Histogram of F_ST_ values for all SNPs; F_ST_ was calculated treating the pelagic species (genera *Rhamphochromis* and *Diplotaxodon*) and benthic species (all other *Malawi* species) as two populations, corresponding to the first split in the radiation (Malinsky et al. 2018). The dashed line shows the top 1% F_ST_ value cutoff. (b) For each of the variant categories described in table 1, the proportion of variants that are in the global top 1% F_ST_ outliers (right of dashed line in panel a) is shown. The third category is expected to be enriched for hybridisation-derived variants. The numbers inside the bars correspond to the number of SNPs in each category. Grey bars correspond to ± 3 block jackknifing standard deviations. (c) For each variant category the proportion of SNPs in different heterozygosity bins is shown. Heterozygosity is averaged across *Malawi* samples. (d) Proportion of F_ST_ outliers in the variant categories described in table 1 stratified by heterozygosity. Error bars correspond to ± 3 block jackknifing standard deviations.

The strong excess of highly differentiated variants among introgression-derived polymorphism suggests that these variants were under selection during the early phases of divergence. However, the excess of high F_ST_ values could also be a consequence of higher than average allele frequencies of introgression-derived variants in the ancestral population. To investigate this possibility, we assessed the distribution of average heterozygosity across *Malawi* for each of the categories, using this measure as a proxy for estimating ancestral allele frequencies (fig. 3c). We found that hybridisation-derived variants (category 3) show elevated heterozygosity compared to the other variant categories (fig. 3c), as expected, but excess divergence was mainly driven by low heterozygosity variants (fig. 3d). Therefore, this result does not appear to be caused by elevated ancestral allele frequencies of introgression-derived variants. Further evidence for this conclusion was obtained from neutral simulations, which showed that such a disproportionate effect on divergence is not expected in the absence of selection (supplementary figs. S6-S8, Supplementary Materials online).

In addition to introgression-derived variants being enriched for outlier loci, we found that these variants show elevated average F_ST_ divergence (supplementary fig. S9) and absolute allele frequency divergence (supplementary fig. S10). This effect is also apparent for the majority of more recent splits in *Malawi* (supplementary fig. S9, S10), suggesting that selection on introgression-derived variants may have played a role at multiple stages of the radiation.

### Long parental haplotypes are fixed in *Malawi* –hybridisation-derived variants are interspersed along the genome

To gain insight into the time scale of mixing of parental haplotypes in *Malawi* (i.e. *Victoria* or *Ruaha Blue* related haplotypes), we considered all fixed differences between the parental lineages, including differences not polymorphic in *Malawi*, and asked over what distance the ancestry states of SNPs in *Malawi* samples are correlated. As expected, the joint probability that two SNPs in a given *Malawi* individual are both homozygous for the *Victoria* allele or both homozygous for *Ruaha Blue* allele decreases with distance between the SNPs (fig. 4a). This correlation in ancestry states extends to several kilobases (kb) (red dots in fig. 4a), which suggests that *Malawi* individuals harbour parental haplotypes several kb in length. Recombination breaks up haplotypes at a rate of one in 30-50 megabases per generation, so this correlation pattern would correspond to haplotype fixation over a few tens of thousands of generations. In contrast to this overall pattern, fixed differences between *Victoria* and *Ruaha Blue* that are polymorphic in Malawi show much shorter range correlation in ancestry states between loci (golden dots in fig. 4a), consistent with them remaining polymorphic for much longer, allowing more time for recombination to break down linkage disequilibrium.

**FIG. 4.**
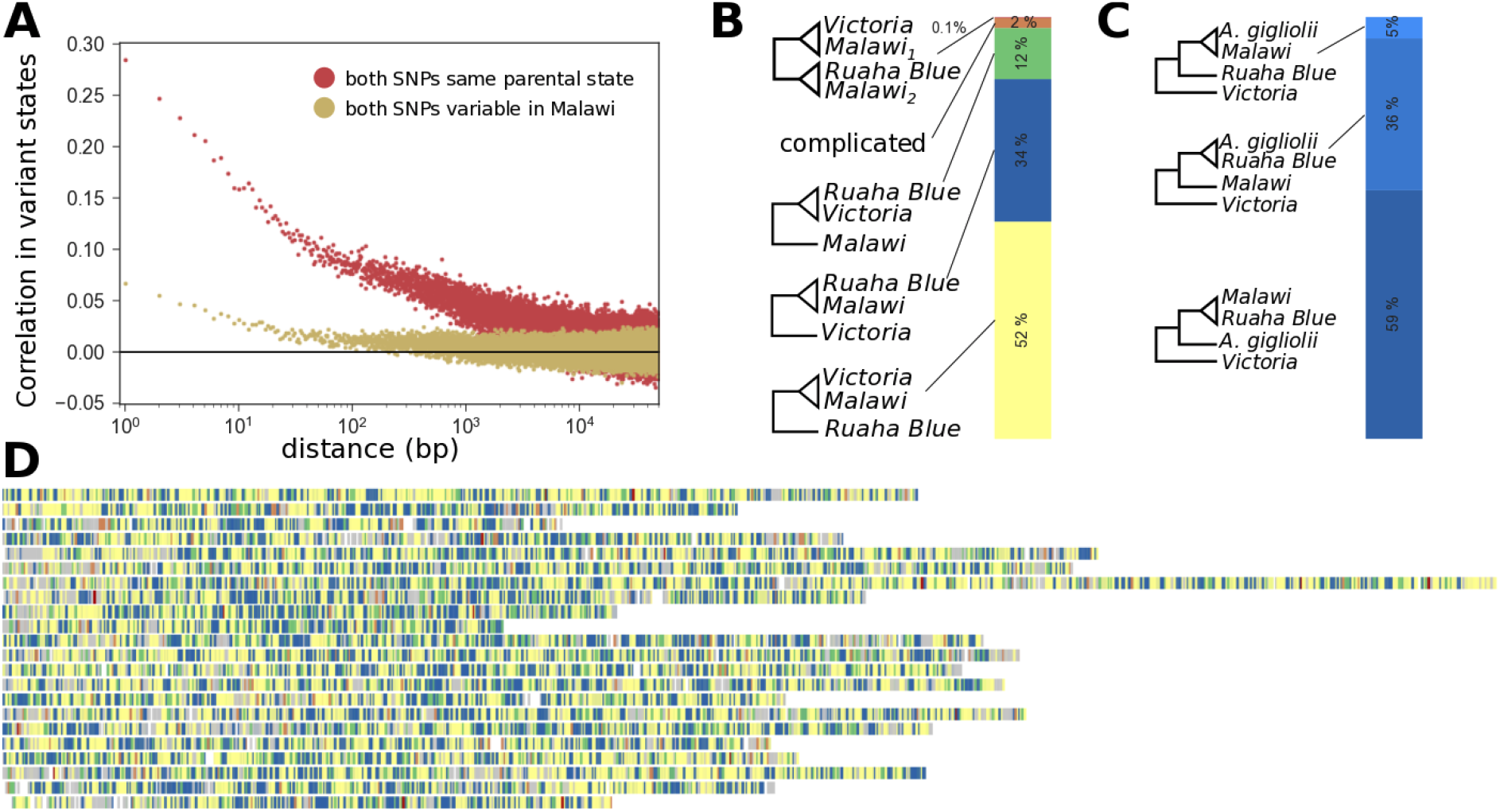
Correlation in variant states and distribution of relatedness patterns along the genome. (a) Genome-averaged excess probability that two SNPs of a certain physical distance that are differentially fixed between *Victoria* and *Ruaha Blue* are of the same state in *Malawi*. Red dots: Joint probability that two variants in *Rhamphochromis woodi* at the given distance are homozygous for the same parental state (*Victoria* or *Ruaha Blue*) minus the expectation if the two variant states were independent (Materials and Methods). Golden dots: Joint probability that two variants at the given separation are variable in *Malawi* minus the expectation if the two variants were independent (Materials and Methods). (b) Neighbour-joining trees of all samples were constructed in overlapping 5kb windows (1kb step-size) and rooted with the Tanganyikan outgroup *Neolamprologus brichardi*. Shallow trees with less than five SNPs separating the common ancestor of *Victoria*, *Malawi*, and *Ruaha Blue* from all the tips of at least two of these groups were excluded (10% of the windows). The remaining trees were classified into five mutually exclusive patterns: (1) All *Malawi* samples are more closely related to all *Victoria* samples than to any *Ruaha Blue* sample (yellow). (2) All *Malawi* samples are more closely related to all *Ruaha Blue* samples than to any *Victoria* sample (blue). (3) All *Victoria* samples are more closely related to all *Ruaha Blue* samples than to any *Malawi* sample (green). (4) Some *Malawi* samples are more closely related to all Victoria samples than to any *Ruaha Blue* sample, while other *Malawi* samples are more closely related to all *Ruaha Blue* samples than to any *Victoria* sample (red, 0.1% of the windows). (6) More complicated relatedness patterns (brown). (c) Sub-classification of pattern (2) into cases (from dark to light blue): all *Malawi* samples are closer to all *Ruaha Blue* samples than to any *A. gigliolii* sample; at least some *A. gigliolii* samples are closer to Ruaha Blue; all *Malawi* samples are closer to all *A. gigliolii* samples than to any *Ruaha Blue* sample. (d) Distribution of relatedness patterns along chromosomes. Regions with ‘shallow trees’ (see above) are shown in grey.

To better understand the genomic distribution of parental haplotypes in *Malawi*, we divided the genome into overlapping chunks of five kb with four kb overlap and constructed NJ trees of all samples for each window (fig. 4b-d). We found that in 52% of the trees all *Malawi* samples clustered with *Victoria*, in 34% of the trees all *Malawi* samples clustered with *Ruaha Blue*, in 12% of the trees *Victoria* and *Ruaha Blue* clustered together and *Malawi* samples formed an outgroup, and the remaining 2% had more complicated topologies (fig. 4b). Notably, only 0.1% of the trees showed a pattern where some *Malawi* samples clustered with *Victoria* while others clustered with *Ruaha Blue*, consistent with long hybridisation-derived haplotypes. This is in contrast to 5.6% of the *Victoria* – *Ruaha Blue* fixed-differences being polymorphic in *Malawi* (that is, in the "hybridisation-derived" category). Furthermore, although the density of hybridisation-derived variants is six times higher in genomic windows in which some *Malawi* samples cluster with *Victoria* and some with *Ruaha Blue* (0.82 per kb compared to 0.11 per kb and 0.15 per kb for windows in which all *Malawi* samples cluster with *Victoria* and *Ruaha Blue*, respectively), more than 99% of the hybridisation-derived variants are outside of these windows. These observations add further evidence that the parental haplotypes carrying hybridisation-derived polymorphisms are generally much shorter than 5 kb.

Overall, there is substantial variation in the topology of relationships between *Malawi*, *Victoria* and *Ruaha Blue* lineages at the 5 kb scale. The observation that *Malawi* clusters with *Ruaha Blue* in 34% of trees, whereas *Victoria* does so only in 12% of trees, supports a substantial contribution from the hybridisation event we have described, but the presence of 12% (*Victoria*, *Ruaha Blue*) trees suggests either an additional component from incomplete lineage sorting (ILS) during the separation of the three groups, or a more complex history involving multiple events. For 59% of the (*Malawi*, *Ruaha Blue*) trees, *Malawi* is closer to *Ruaha Blue* than *A. gigliolii* (fig. 4c), while only for 5% of these trees the opposite is true, consistent with the proposed hybridisation event in the presence of some ILS in the ancestry of the *Ruaha Blue/A. gigliolii/Malawi* contributor lineages. Together, this suggests that the genomes of *Malawi* samples are composed predominantly of multi-kilobase haplotypes deriving from *Victoria* or *Ruaha Blue* lineages. The genomic distribution of these haplotypes is non-uniform and often spans across many 5kb windows (fig. 4d), suggesting that at least some regions fixed quite rapidly after hybridisation.

### Hybridisation-derived variants are enriched in vision- and immune defence-related genes

To investigate whether hybridisation-derived variants contribute to specific biological functions, we tested for enrichment of hybridisation-derived variants (category 3 in table 1) in genes annotated by different gene ontology terms (GOs). In particular, we computed the density of such variants across the exons of all genes (normalised by the number of private *Malawi* variants, category 1 in table 1) in each GO category and calculated an empirical enrichment p value (Materials and Methods). The GO categories with the most significant enrichment are related to non-coding and ribosomal RNA functions (*ribonucleoprotein complex biogenesis* p = 0.0001), vision (*visual perception* p = 0.0046) and defence response (*defence response to bacterium* p = 0.005) (fig. 5). Both genes related to vision and to pathogen defence have previously been suggested to play a role in speciation (Carleton et al. 2005; Malmstrøm et al. 2016; Egger et al. 2017; Maan et al. 2017; Meier et al. 2017; Malinsky et al. 2018). Consistent with the results above, we find that even genes in these categories do not show long hybridisation-derived haplotypes, that is, most fixed differences between *Ruaha Blue* and *Victoria* in these regions are not variable in *Malawi*. However, for hybridisation derived variants there is considerable correlation of which *Malawi* individuals have *Victoria* and which have *Ruaha Blue* alleles within a gene (supplementary fig. S11). We obtain similar enrichments when normalising hybridisation-derived variant density by the number of variants shared with a parental lineage (category 2 in table 1; supplementary fig. S12), or by accessible exon length (supplementary fig. S13). In particular, in both cases *visual perception* is also among the most enriched GO terms (p = 0.0006 and 0.0101, respectively).

**FIG. 5.**
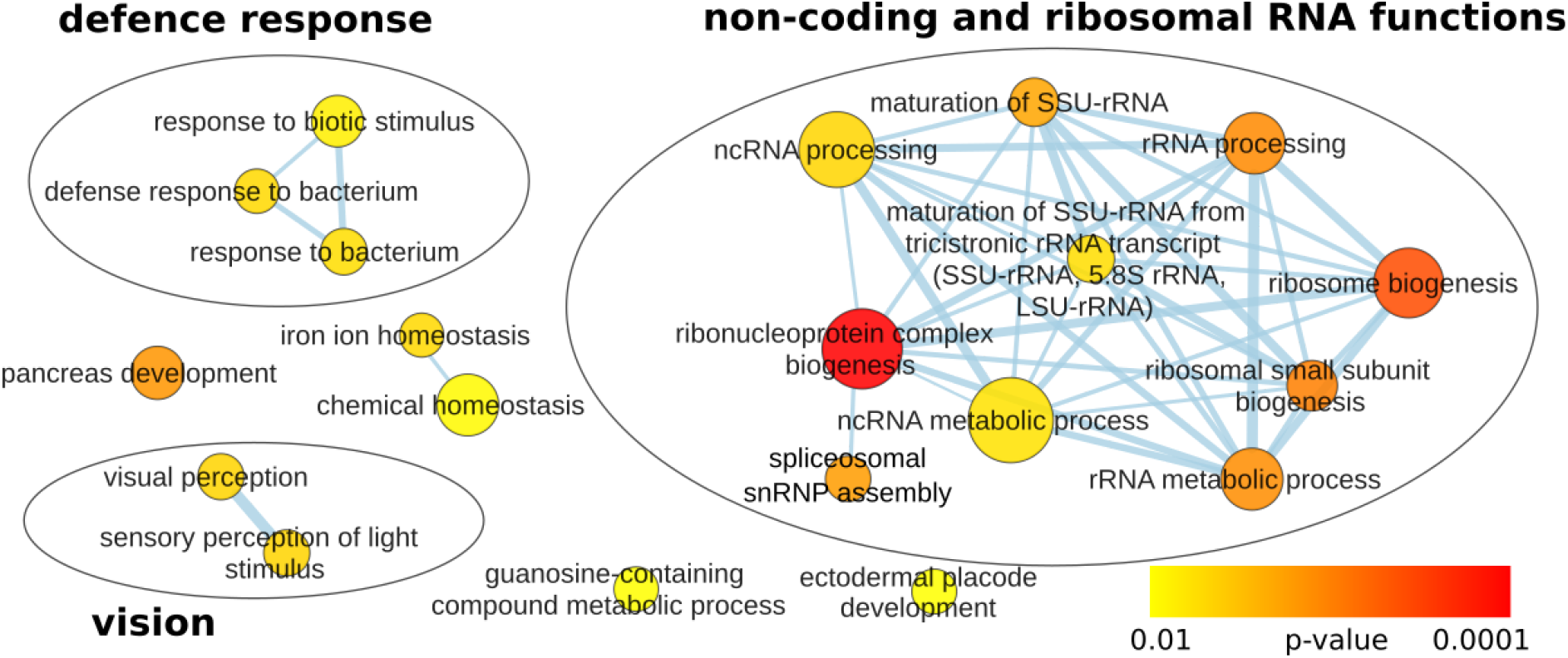
Enrichment of hybridisation-derived variants in biological process gene ontologies (GOs). Genes in the *Oreochromis niloticus* gene annotation were linked to zebrafish GO categories and enrichment of *Malawi* variants differentially fixed between *Victoria* and *Ruaha Blue* (category 3 in table 1) relative to private *Malawi* variants (category 1 in table 1) in exonic regions was tested (Materials and Methods). Categories with enrichment p-values <0.01 are shown. Size of the circles is proportional to the number of genes present in the annotation in each category (from 10 to 220). Analogous analyses with different normalisations are given in supplementary figs. S12 and S13.

### Hybridisation-derived variants assort non-randomly in the first split in the Lake Malawi cichlid radiation

When looking at all variants, we saw in fig. 2c there is not much difference in the relative amounts of *Victoria* and *Ruaha Blue* ancestries in different *Malawi* groups. Consistent with this, there is very little differential contribution of the parental lineages to *benthic* and *pelagic* (f_4_(P_1_=*pelagic*, P_2_=*benthic*, P_3_=*Victoria*, P_4_=*Ruaha Blue*) = 0.08%, block-jackknifing p-value=3.72 ×10^−6^). However, for hybridisation-derived segregating variants there is a clear excess of *Victoria* alleles in *benthic* (f_4_ = 1.43%, p =1.42 ×10^−6^). Interestingly, there is a strong positive association between allele frequency differentiation across the *benthic*/*pelagic* split and excess *Victoria* ancestry in benthic species (supplementary fig. S14). For hybridisation-derived variants with less than 10% absolute allele frequency differentiation between *benthic* and *pelagic*, there is actually a significant depletion of *Victoria* alleles in *benthic* (that is, an excess of *Ruaha Blue*). However, this pattern reverses for more differentiated variants, with hybridisation-derived variants with above 60% allele frequency difference across the first *Malawi* split showing 11% excess *Victoria* ancestry in *benthic* compared to *pelagic* (supplementary fig. S14).

## Discussion

A clear message from the increasing number of genomic investigations of non-model organisms is that hybridisation between related species is far more common than previously thought, leading to fundamentally reticulate patterns of evolution, where species relationships are represented by networks rather than trees. In many cases it has been shown that this process can transport selectively favourable alleles between species and thus contribute to adaptation (Herman et al. 2018; Suarez-Gonzalez et al. 2018; Walsh et al. 2018; Moest et al. 2019). However, the effect of such hybridisation events on biological diversification, that is, whether it increases or reduces the rate at which new species emerge over time is not generally understood. While empirical evidence for a direct role of hybridisation in speciation events that do not involve polyploidization remains relatively rare (Schumer et al. 2014), a number of recent genomic studies suggest that hybridisation of genetically divergent lineages can allow for the reassembly of old genetic variation and thereby play a decisive role in rapid adaptive diversification (reviewed in Marques et al. 2019).

Meier et al. (2017) were the first to establish a functional role of ancestral hybridisation in an adaptive radiation, by showing that hybridisation-derived variation was under selection during species divergence in the Lake Victoria Region Superflock of cichlids. Here we demonstrate that the Lake Malawi cichlid adaptive radiation was also seeded by a hybridisation event providing genetic variation that was subsequently differentially selected during divergence of the major eco-morphological clades. In particular, variants that are differentially fixed between the two parental lineages show striking excess divergence across the benthic/pelagic split of Malawi cichlids and also across more recent splits. Together with recent data from Lake Tanganyika cichlids (Irisarri et al. 2018), where hybridisation at the base of radiating lineages has also been described, although not yet demonstrated to be functionally important, this suggests that hybridisation might have repeatedly played an important role in the remarkable patterns of adaptive speciation in East African cichlids.

A hybrid origin of the Lake Malawi cichlid adaptive radiation had previously been proposed by Joyce et al. (2011) on the basis of mitochondrial-nuclear discordance. In particular, they suggested that two divergent riverine *Astatotilapia calliptera*-like lineages seeded the species rich Mbuna clade of Malawi cichlids. However, our recent genome-wide investigation including geographically broad sampling of *A. calliptera* has clearly shown that all *A. calliptera* lineages are nested within the Lake Malawi cichlid radiation (Malinsky et al. 2018). In contrast, the lineage we found contributing to the Malawi ancestral population – *A.* sp. ‘ruaha blue’ – is a genome-wide outgroup to both the Lake Malawi and Lake Victoria radiations. This means that the hybridisation event described here is much older than the one described by Meier et al. (2017) both in terms of the date of the event itself and in terms of the divergence between the parental lineages. Indeed, both of the parental lineages of the LVRS identified in Meier et al. (2017) are contained in our *Victoria* clade (fig. 1b, top clade).

It is notable that the present-day habitat of the only known descendant of one of the parental lineages of the Lake Malawi cichlid radiation, *A.* sp. "ruaha blue", is adjacent to that of the Lake Malawi radiation, but separated by the high-altitude Livingstone/Kipengere mountain range estimated to have formed during the Pliocene rifting that formed Lake Malawi. However, fossil evidence supports a connection between the Ruaha/Rufiji system and the Lake Malawi basin 2-3.75 million years ago (Stewart and Murray 2013), consistent with our timings of the separations of the relevant lineages.

Characterising the distribution of parental ancestry along Malawi cichlid linkage groups, we found that the genomes of Malawi cichlids are mosaics of *Victoria-* and *Ruaha Blue*-like ancestry (the latter making up ∼34%). *Ruaha Blue*-like haplotypes extend over several kbs suggesting that most polymorphism resulting from the hybridisation event was fixed relatively quickly after the hybridisation event in one or the other direction, within tens of thousands of years. Furthermore, there is a clear signal of non-random distribution of the different ancestries across the genome, which is consistent with recent observations in Princess cichlids from Lake Tanganyika (Gante et al. 2016). This might be the result of variation in local recombination rate, which has been shown to affect rates of introgression (Martin and Jiggins 2017). A high-resolution recombination map for Malawi cichlids will allow the testing of the effect of recombination rate variation on ancestry distribution in the future.

In the majority of cases all *Malawi* samples have the same parental haplotype in a given genomic region. At first sight this seems inconsistent with the "combinatorial view on speciation and adaptive radiation" suggested by Marques et al. (2019), but it is consistent with results from the LVRS, where there is a strong correlation of parental ancestries across species (Meier et al. 2017). In Malawi cichlids, hybridisation-derived segregating variants are relatively rare and interspersed amongst the longer, fixed parental haplotypes. A possible explanation for this pattern is that some variants managed to escape the initial fixation process through recombination or gene conversion, possibly because they were under balancing selection across a meta-population of diverging eco-types. This would suggest that much of the hybridisation derived variation that has been maintained is functional, which could explain why, despite this ancestral hybridisation event, Malawi cichlid lineages show overall remarkably little genetic diversity (Malinsky et al. 2018).

We further show that hybridisation-derived variants are enriched in genes related to vision, pathogen defence and non-coding and ribosomal RNA function. The former two categories have previously been implicated to play a role in speciation and adaptive radiation in cichlids or other fishes (Carleton et al. 2005; Malmstrøm et al. 2016; Maan et al. 2017; Meier et al. 2017; Malinsky et al. 2018). In particular, Meier et al. (2017) have shown that in the LVRS divergent parental haplotypes of a short-wavelength opsin gene differentiate clear shallow water algivores from deep turbid water detritivores. In our case, while several vision-related genes carry hybridisation-derived variants, there is no separation of Malawi cichlids into species carrying *Victoria-like* and *Ruaha-Blue*-like haplotypes along a whole gene-body. This is consistent with our observation that hybridisation-derived polymorphism is not located on long haplotypes. The observation of long-hybridisation derived haplotypes in Meier et al. (2017) is restricted to a single gene. A comparison of whole-genome sequencing data from these two independent instances of ancestral hybridisation prior to radiation would allow more general understanding of the segregation patterns of parental haplotypes.

The enrichment of hybridisation-derived genetic variants in non-coding RNA and ribosome-related gene ontologies was unexpected. While we applied conservative filtering to genetic variants (see Materials and Methods), we cannot totally exclude the possibility that this pattern is related to the repetitive nature of many of the genes in these categories. However, if this pattern were due to sequencing reads from repetitive regions collapsed to the same reference position ("para-SNPs"), then there is no reason that this pattern would be specific to hybridisation-derived variants. On the contrary, we would rather expect enrichment for genetic variants that are also polymorphic in the parental lineages. A biological explanation for the enrichment of hybridisation-derived variants in non-coding-RNA-related genes might be connected to differential control of transposable elements (TEs) in the parental lineages (Wheeler 2013). Indeed, there is accumulating evidence that TEs play a role in reproductive isolation and speciation (Serrato-Capuchina and Matute 2018). This could be addressed by a thorough investigation of TE activity and distribution across Malawi cichlids and parental lineages and in crosses.

While overall across the whole genome there is little difference in the relative contributions of parental lineages to different *Malawi* lineages, this is not true for hybridisation-derived variants. For hybridisation-derived variants that are highly differentiated between pelagic and benthic *Malawi* lineages (and thus expected to have been under divergent selection) it is significantly more likely to find the *Victoria* allele in benthic rather than in pelagic species. This signal could be driven by divergent ecological selection, for example, if the two parental lineages had different "pre-adaptations" to the different niches, possibly reflecting experience of different habitats prior to introgression. This fits with the idea that introgression brings added potential by mixing up adaptive genomic combinations of the introgressing lineages (Marques et al. 2019). An alternative explanation is selection to sort out genetic incompatibilities between alleles at different loci that accumulated during divergence of the parental lineages (Stelkens et al. 2010). However, given the above observation that most parental haplotypes are actually fixed one way or the other, it is not clear whether one would expect the remaining hybridisation-derived variants to show incompatibilities. We suggest that the direct investigation of present-day hybrid populations and crosses will be necessary to tease these effects apart and map genetic incompatibilities.

In this study, we infer a hybrid origin of the ancestral population of the Malawi cichlid adaptive radiation. We demonstrate that while much of the hybridisation-derived variation has apparently been quickly fixed for one or the other parental haplotype in the *Malawi* ancestor, the variation that was maintained contributes more to early differentiation among Malawi cichlid lineages than expected in the absence of selection. While we present the geographically most comprehensive whole-genome dataset of haplochromine cichlids to date, the number of samples per species/population is mostly restricted to a single diploid individual. More comprehensive population sampling of parental lineages and species of the Lake Malawi cichlid adaptive radiation will enable more detailed characterisation of hybridisation-derived variants and their evolutionary significance.

Finally, it is worth noting that the key lineage in uncovering the evidence for introgression in the ancestor of the Malawi radiation, *A.* ‘ruaha blue’, appears to be represented by a single extant species confined to part of a single river system geographically remote from Lake Malawi, and entirely undocumented prior to collection of the samples presented here in 2012. Had this lineage gone extinct or not been sampled, we would have been missing direct evidence for the hybrid origin of the Malawi haplochromine lineage. This suggests that there may be ample opportunity for false negatives in the study of the influence of hybridisation on adaptive radiation. Judging from our results and other recent findings in cichlid fishes and beyond, it is not impossible that hybridisation indeed is the dark matter (or fuel) of rapid ecological diversification.

## Materials and Methods

### Sampling

Sequencing reads for some samples were obtained from previous studies (Brawand et al. 2014; McGee et al. 2016; Malinsky et al. 2018). For newly sequenced samples, ethanol-preserved fin-clips were collected by M. J. Genner, A. Indermaur, A. Lamboj, B. Ngatunga, F. Ronco, W. Salzburger, and G. F. Turner between 2004 and 2014 from Malawi, Tanzania, and Zambia, in collaboration with the Fisheries Research Unit of the Government of Malawi (various collaborative projects), the Tanzania Fisheries Research Institute (MolEcoFish Project), and the Department of Fisheries, Republic of Zambia (see supplementary table 1 for sample details and NCBI accession numbers). Samples were collected and exported with the permission of the Fisheries Research Unit of the Government of Malawi, the Tanzania Commission for Science and Technology, the Tanzania Fisheries Research Institute, and the Department of Fisheries, Republic of Zambia.

### Sequencing, variant detection and filtering

New samples were sequenced on an Illumina HiSeq platform (100–125 bp paired-end reads) to a fold coverage of ∼15X (supplementary table 1). All samples were aligned to the tilapia reference genome Orenil 1.1 (GenBank assembly accession: GCA_000188235.2) using bwa-mem (Li 2013). Duplicate reads were marked on both per-lane and per sample basis using the MarkDuplicates tool from the Picard software package with default options (http://broadinstitute.github.io/picard). Local realignment around indels was performed on both per lane and per sample basis using theIndelRealigner tool from the GATK v3.3.0 software package. Per-sample variant detection was performed with GATK v.3.5 HaplotypeCaller (McKenna et al. 2010) and joint genotyping then performed using the GenotypeGVCFs tool. Because the dataset contains samples from many species, we set a flat per-locus prior likelihoods using the --input_prior option. At this stage we also included non-variant sites.

Because of mapping to a relatively distant reference (∼3% divergence in our filtered dataset and ∼6% over fourfold degenerate sites in Brawand et al. (2014)), we used more stringent filtering criteria than in previous studies (Malinsky et al., 2015, Malinsky et al., 2018). First, we masked out all sites in the reference where any overlapping 50-mers (sub-sequences of length 50) could not be matched back uniquely and without 1-difference. For this we used Heng Li’s SNPable tool (http://lh3lh3.users.sourceforge.net/snpable.shtml), dividing the reference genome into overlapping 50-mers, and then aligning the extracted 50-mers back to the genome (we used bwa aln -R 1000000 -O 3 - E 3). Next, using the all site VCF, we masked out any sites where more than 10% of mapped reads had mapping quality zero, or where the overall mapping quality (MQ) was less than 40, sites where the number of no-called samples (NCC) was greater than one, sites which were within 3bp of indels in any sample, and sites where the sum of overall depth for all samples was unusually high or low, with the depth cutoffs corresponding to ∼22% and ∼98% percentiles of the distribution (supplementary fig. S15). This resulted in 338 Mb of "accessible genome" on the 22 linkage groups (not including unplaced scaffolds), about half of the total placed genome size of 657 Mb. The VCF was then subset to include only SNPs, to which additional filters were applied for excess heterozygosity (InbreedingCoeff < −0.5, ExcessHet >=20) and low quality by depth (DP > 2) resulting in a final dataset of 18,760,777 SNPs.

### Patterson’s D and f4 admixture ratio

We calculated Patterson’s D and f4 admixture ratio statistics as defined in (Patterson et al. 2012) using custom scripts available at https://github.com/feilchenfeldt/pypopgen. Calculations were performed on all SNPs that passed filtering and block-jackknife standard-deviations were calculated across chromosomes (Green et al. 2010).

### Divergence statistics

*Malawi* samples were split into pelagic and benthic species, corresponding to the first split in the phylogenetic tree in Malawi (pelagic samples: *Rhamphochromis woodi*, *R. logiceps*, *Diplotaxodon* sp. ‘macrops ngulube’, *D. limnothrissa*). For all passed SNPs that were segregating within *Malawi*, the fixation index, F_ST_, was calculated between pelagic and benthic species using the Weir Cockerham estimator implemented in VCFtools 0.1.14 (Danecek et al. 2011) (option --weir-fst-pop). Absolute allele frequency differentiation of monophyletic subgroups of Malawi samples, A and B, was calculated as ABS(allele frequency A - allele frequency B) (supplementary fig. S10).

### Genomic distribution of parental ancestry

We considered all variants that were differentially fixed between *Victoria* and *Ruaha Blue* (regardless of whether they were variable in *Malawi* or not) and recorded for each of these variants (1) the state in a given focal *Malawi* individual (either homozygous for *Victoria* allele, heterozygous, or homozygous for *Ruaha Blue* allele) and (2) whether they were variable in *Malawi* or not. Then, we considered all pairs of variants of a given genomic distance in bps (only variant pairs with distance < 50 kb were considered), and calculated the genome-wide fraction of variant pairs of the same state (e.g., both homozygous for the *Victoria* allele in the focal individual, both variable in *Malawi*, etc.) and the genome-wide single variant probability for each of the states. These statistics were used to calculate the empirical excess probability of observing two variants of a given distance in the same state relative to observing any two variants in the same state. In particular, the red dots in fig. 4a correspond to 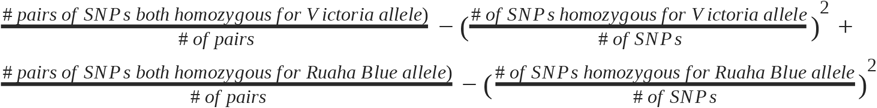 where the focal individual was chosen as *Rhamphochromis woodi*. Results for different focal individuals are indistinguishable from the ones in fig. 4a. Analogously, the golden dots in fig. 4a correspond to 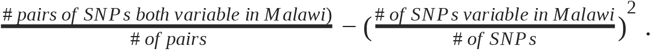

Next we split the genome into overlapping windows of 5kb with 1kb offset. For each window, pairwise differences between all samples were calculated using custom scripts available at https://github.com/feilchenfeldt/pypopgen and neighbour-joining (NJ) trees were constructed using Biopython’s Phylo package. We excluded shallow trees with fewer than five SNPs separating the common ancestor of *Victoria*, *Malawi*, and *Ruaha Blue* from all the tips of at least two of these groups from further analysis (10% of the windows). This approach was chosen to remove genomic regions with shallow, uninformative trees. The remaining trees were classified according to their topology into the four patterns as described in the caption of fig. 4.

### Gene enrichment test

We downloaded the gene annotation for Orenil 1.1. release 102 from the NCBI website, and retained for further analysis the 9452 genes located on the 22 linkage groups that were annotated with a HGNC gene symbol. Using Bioconductor 3.5 in R 3.5.1, we downloaded zebrafish GO annotations (org.Dr.eg.db) for the category biological process, compiled a mapping between GO categories and gene symbols present in Orenil 1.1 102 and retained the 1641 GO categories with between 10 and 1000 genes. For each gene, we computed the number of exonic variants of each of the four categories defined in table 1. Splice variants were ignored for this analysis (i.e., all exons were collapsed). We also calculated the number of accessible sites (see above) for each of the exonic regions. Next, we calculated for each GO category the total number of exonic gene variants in each of the four variant categories as well as the corresponding accessible genome lengths and computed densities of hybridisation-derived variants using the following normalisations (c.f., table 1): (1) (# of category 3 variants)/(# of category 1 variants), (2) (# of category 3 variants)/(# of category 2 variants), (3) (# of category 3 variants)/(accessible genome length). To test whether these variant densities are significantly higher than expected, we computed an empirical background distribution of variant densities for randomly assembled GO categories of the same number of genes and calculated empirical p-values as 1-RANK/10000, where RANK is the rank of the variant density of a GO category among the background distribution values and where 10000 background observations were computed for each observed category size.

For each normalisation, the overlap of GO categories with enrichment p-values < 0.01 were visualised using the Cytoscape 3.7.1 Enrichment Map App using an edge cutoff of 0.375.

## Supplementary Material online

Available as separate files.

## Acknowledgements

We thank A. Indermaur, A. Lamboj, F. Ronco and the staff of the Tanzania Fisheries Research Institute for help with sample collection. We acknowledge funding from Wellcome Trust grants WT206194 and WT207492 (H.S. and R.D.), the European Research Council, ERC CoG “CICHLID∼X” (617585) and Swiss National Science Foundation, grant nr. 176039 (W.S) and the Royal Society – Leverhulme Trust Africa Awards AA100023 and AA130107 to MJG, BPN and GFT. Sequencing reads used in this study are submitted to the NCBI short read archive (see supplementary table 1 for accession numbers) and variant call (VCF) data from this study is available from the European Nucleotide Archive accession number XYZ.

