## Supplementary material for "Ancestral hybridisation facilitated species diversification in the Lake Malawi cichlid fish adaptive radiation": Supplmentary Online Material

#### Simulations

We performed neutral coalescent simulations using msprime (Kelleher et al. 2016) to test whether the observed excess-divergence of hybridisation-derived variation is observed under neutral evolution. The simulations were designed to qualitatively match the diversity patterns observed in the real data (supplementary fig. S1).

In particular, within-sample effective population sizes were inferred from real data as $N_e = \pi/(4\mu)$, where $\pi$ is the number of pairwise genetic differences per accessible site in the genome, and $\mu$ is the per generation mutation rate set to 3*3.5e-9 as inferred in (Malinsky et al. 2018) and assuming a generation time of three years. Since alignment here was performed against the relatively distant outgroup genome Orenil. 1.1 (Tilapia) with an average pairwise sequence divergence of ~6% over fourfold degenerate sites in Brawand et al. (2014), and conservative genome filtering was performed (see Materials and Methods), there is a certain bias in our accessible genome to overall more conserved genomic regions. Consistent with this, divergence to the reference genome in our accessible genome is only ~3%. To account for this, we scaled all branch length estimates with a scaling factor of 1.53, which scales the date of the split between pelagic (*Rhamphochromis*, *Diplotaxodon*) and other Lake Malawi cichlid species to the 460 thousand years estimated by Malinsky et al. (2018).

Pairwise split times of two lineages *x* and *y* were estimated as

$$T_{split} = \frac{d_{xy} - (\pi_x + \pi_y)/2}{2\mu}, \qquad (1)$$

where $d_{xy}$ is cross-lineage pairwise sequence divergence per accessible site. A split tree was constructed from these data using the neighbour-joining algorithm. The unequal length of terminal branches, which could be the result of variation in mutation rates, generation time, or drift was implemented in the simulations by setting time zero for the longest branch and treating shorter branches as ancient samples (times given in fig. S1).

Effective population sizes on internal branches were set to the average of the two child branches. Note that for computational efficiency, in the simulations the non-Malawi lineages *Victoria*, *Ruaha Blue*, *A. gigliolii*, and *H. vanheusdeni* were each represented by panmictic (meta)populations with effective population sizes set to match the overall diversity patterns in these (inhomogeneous) groups. For these branches, when calculating $N_e$ of parental branches, the average of the actual sample effective population sizes was used rather than the $N_e$ of the metapopulation.

Two instantaneous gene flow events were added to the simulations: Gene flow into the common ancestor of the Malawi radiation from the *Ruaha Blue* branch as inferred in fig. 2, and, within *Malawi*, gene flow from the pelagic clade *Diplotaxodon* into the benthic clade, which constitutes the strongest gene flow event among major *Malawi* groups inferred in Malinksy et al. (2018). Note that simulation input gene flow fractions of 22% and 31%, respectively, are set to match inferred $f_4$ admixture fractions of 10% each. Furthermore, we lengthened the branch of the ancestor of the non-pelagic Malawi clade by 10k generations, because an expected effect of the observed gene flow is to shorten this branch, and otherwise benthic species cluster with the pelagics in the simulation results.

The simulation results give a reasonable match to the data (supplementary figs. S2, S6, S7). Cross-species pairwise differences and $F_{ST}$ are slightly higher in the simulations than in the real data. This can be explained by additional gene flow events between Malawi species that have not been modelled here (see Malinsky et al. 2018) leading to larger amounts of shared variation between non-sister-branches.

In the neutral simulations, the proportion of $F_{ST}$ outliers is elevated for both old shared variation and hybridisation-derived variation relative to private *Malawi* variants for low heterozygosity bins (fig. S8, right panel). But importantly, there is no excess divergence of hybridisation-derived variants as compared to old shared variation. For the heterozygosity bins 0.2-0.3 and higher, there is no significant difference between any of the variant categories. The difference between private and non-private variation in the heterozygosity bins 0-0.1 and 0.1-0.2 is due to the fact that non-private variants are relatively enriched at the right end of these bins.

### Supplementary Figures

---

**FIG. S1**

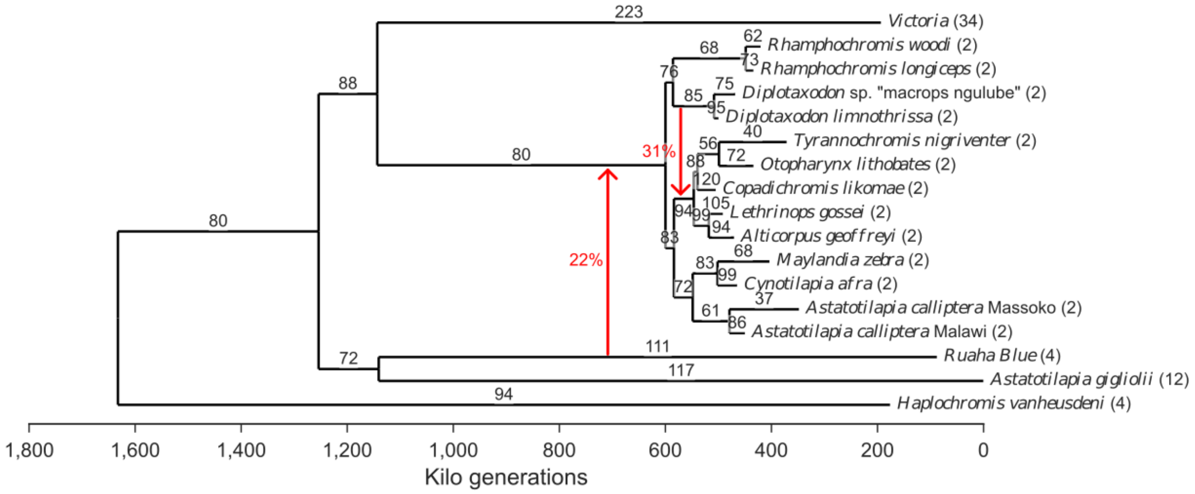


Input species split tree for the coalescent simulations in msprime (Kelleher et al. 2016). Numbers on branches indicate piecewise-constant effective population sizes in thousands. Red arrows and associated percentages indicate gene-flow events ("MassMigrations" in msprime). Numbers in parenthesis right of sample names indicate haploid sample sizes.

---

**FIG. S2**

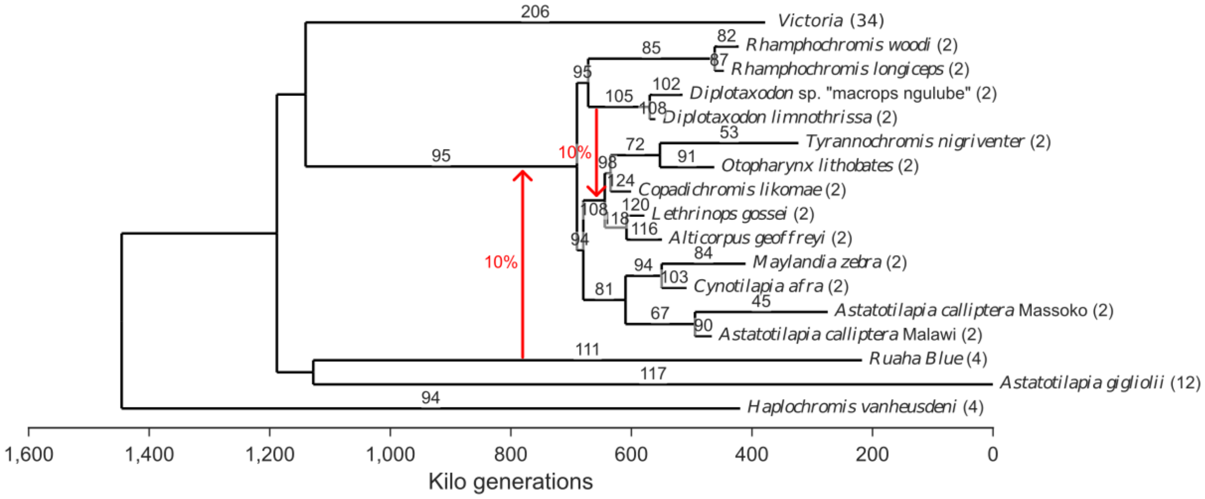


Species tree inferred from the coalescent simulations (see fig. S1 for simulation input). Numbers on branches indicate piecewise-constant effective population sizes. Red arrows and associated percentages indicate gene-flow proportions inferred using the $f_4$ admixture ratio. Numbers in parenthesis right of sample names indicate haploid sample sizes. Note that the split time of the outgroup *H. vanheusdeni* cannot be inferred from the data and is set posteriori.

**FIG. S3**

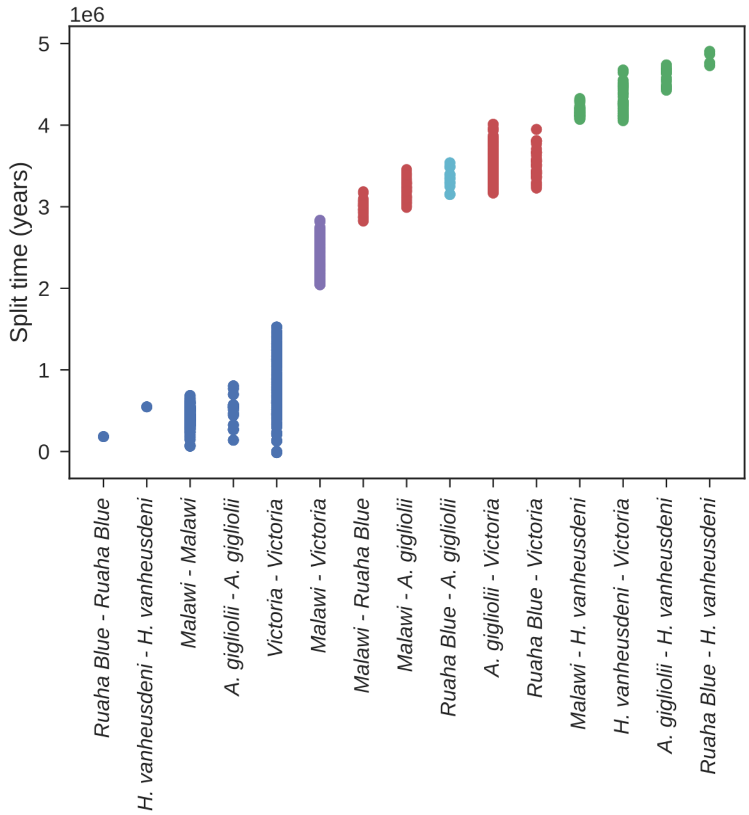


Pairwise sample split time estimates using equation (1), binned by clade, using a mutation rate of 3.5e-9 per bp per year (Malinsky et al. 2018) and a generation time of 3 years. Each dot corresponds to a point estimate obtained from a given pair of individual samples of the two clades on the x-axis. Blue dots correspond to within-group split time estimates. Other estimates are shown in the same colour if they correspond to the same split in fig. 1. Split time estimates assume the absence of gene flow between lineages. Note that mutation rate estimates are subject to uncertainty (Malinsky et al. 2018).

---

**FIG. S4**

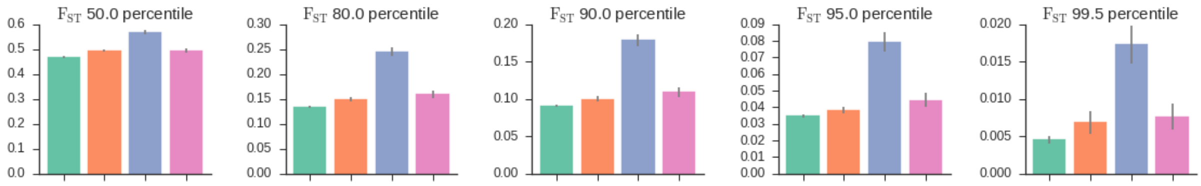


Proportion of $F_{ST}$ outliers in the different SNP categories for different overall percentiles of $F_{ST}$ outliers on real data. The top 99 percentile is shown in fig 3b.

---

**FIG. S5**

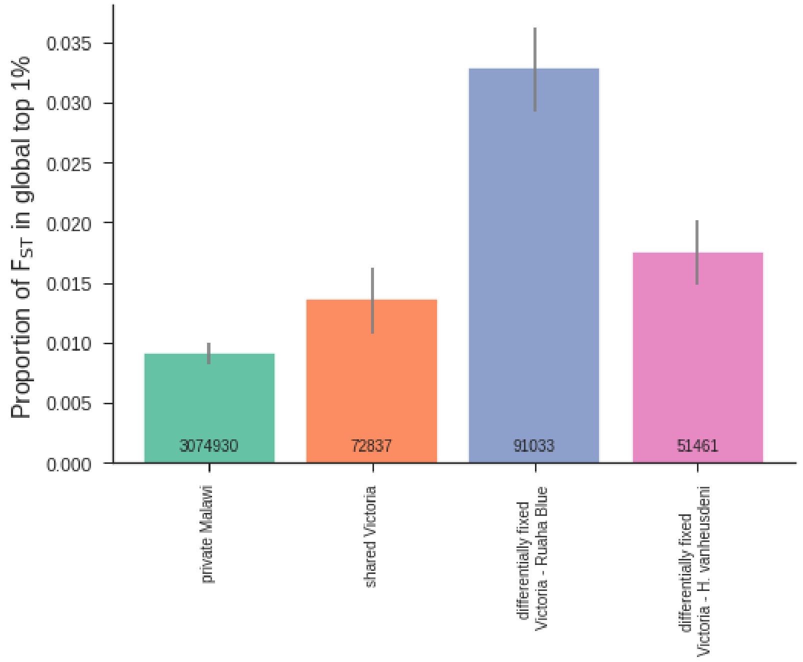


Distribution of $F_{ST}$ outliers across variant categories described in Table 1. This plot is equivalent to fig. 3b, except that the *Victoria* category was subset to two samples, *Paralabidochromis flavus* and *Pundamilia nyererei*, so that it has the same sample size as *Ruaha Blue*.

---

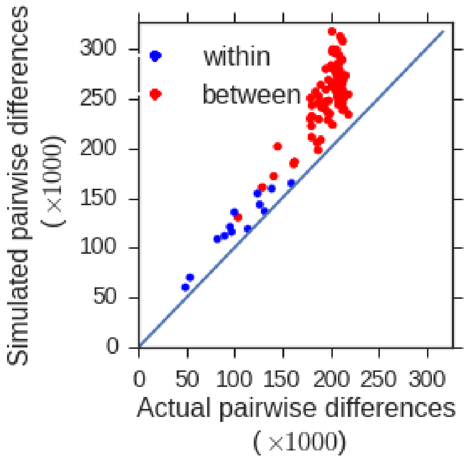


**FIG. S6**

Comparison of pairwise differences between *Malawi* samples in the real data (x-axis) and simulations (y-axis).

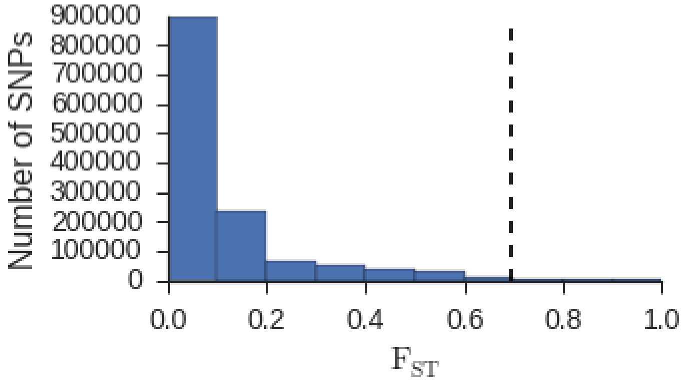


**FIG. S7**

Histogram of $F_{ST}$ values for the pelagic-benthic split inferred from simulated data.

**FIG. S8**

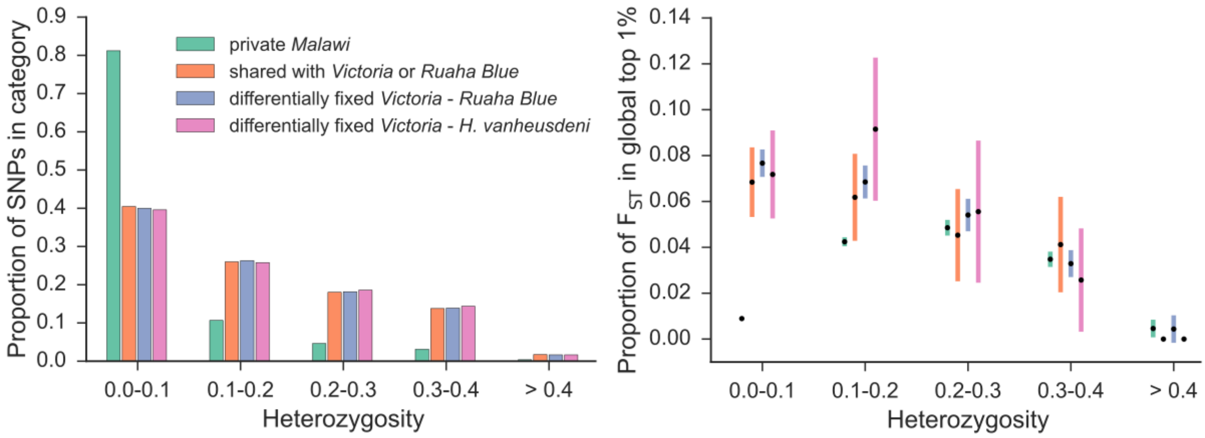


Distribution of $F_{ST}$ outliers across variant categories on simulated data. For each variant category the proportion of SNPs in different heterozygosity bins is shown (left panel) . Heterozygosity is averaged across Malawi samples. Proportion of $F_{ST}$ outliers in the variant categories described in table 1 stratified by heterozygosity (right panel). Error bars correspond to ± 3 block jackknifing standard deviations.

**FIG. S9**

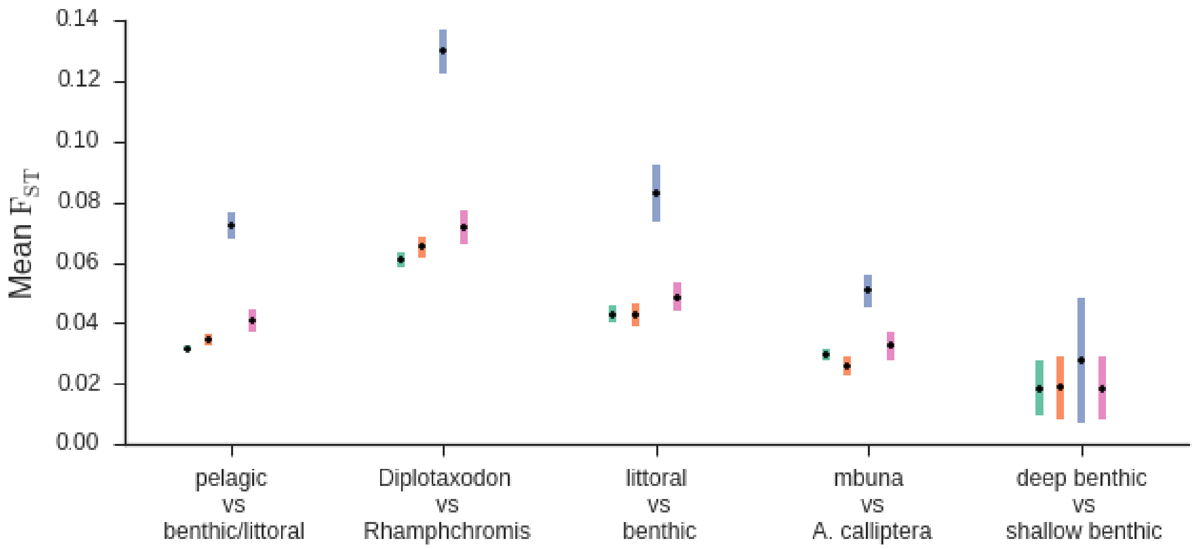


Mean $F_{ST}$ in polymorphism categories for different splits between malawi eco-morphological groups. Color code as in fig. 3. Bars correspond to +/- three block jackknifing standard deviations.

---

**FIG. S10**

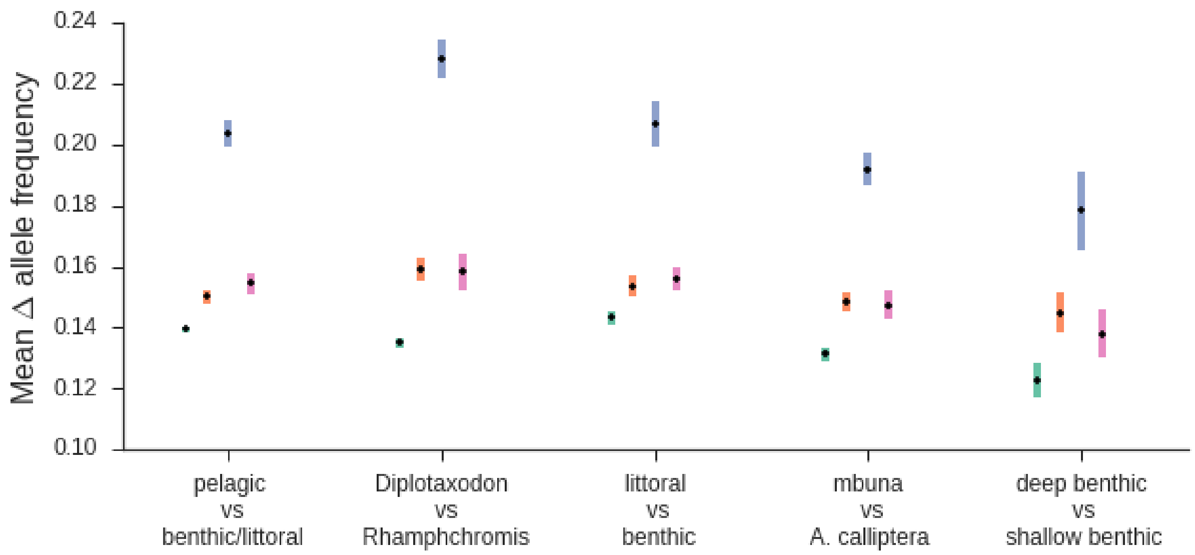


Mean absolute allele frequency difference in polymorphism categories for different splits between malawi eco-morphological groups. Color code as in fig. 3. Bars correspond to +/- three block jackknifing standard deviations.

---

**FIG. S11**

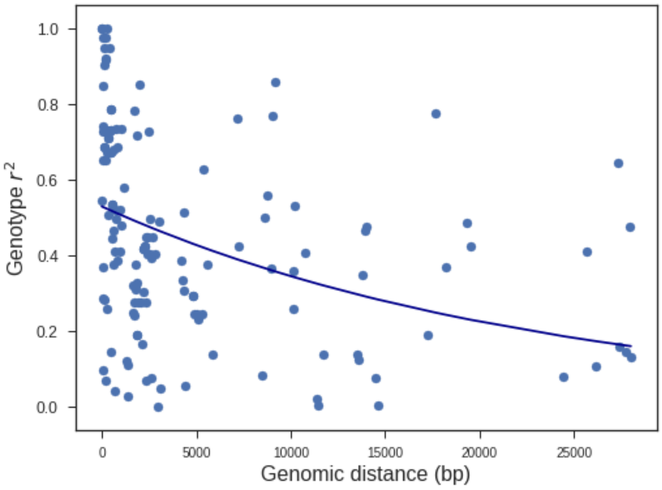


Genotype correlation of pairs hybridisation-derived variants in genes of the vision GO category as a function of genomic distance. Genotype correlation was calculated using the --geno-r2 option in vcftools (Danecek et al. 2011).

---

**FIG. S12**

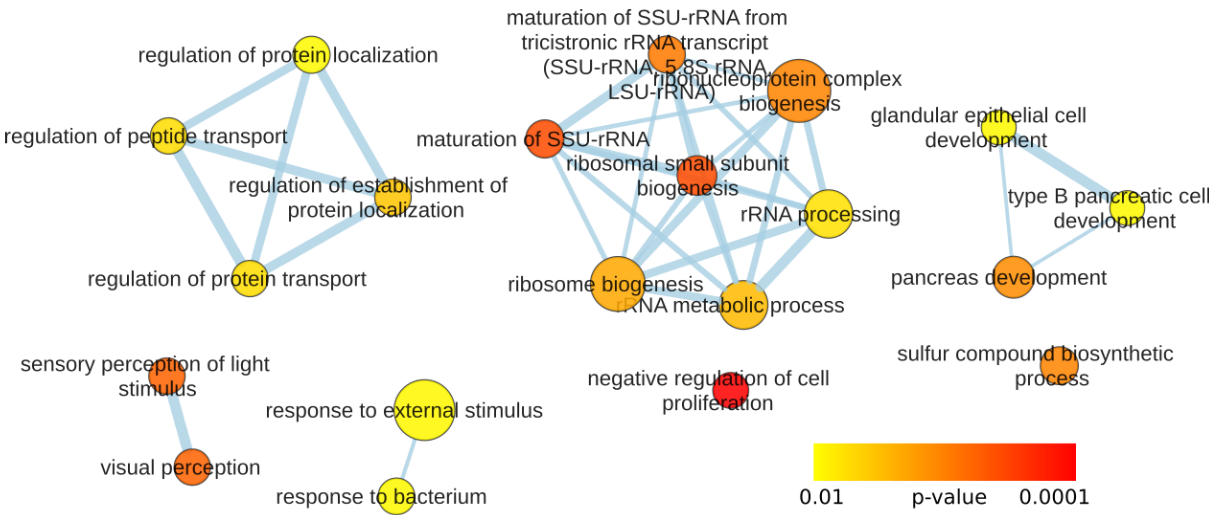


Enrichment of hybridisation-derived variants in biological process gene ontologies (GOs). Genes in the *Oreochromis niloticus* gene annotation were linked to zebrafish GO categories and enrichment of *Malawi* variants differentially fixed between *Victoria* and *Ruaha Blue* (category 3 in table 1) relative to variants that are polymorphic in both Malawi and a parental lineage (category 2 in table 1) in exonic regions was tested (see Materials and Methods). Categories with enrichment p-values <0.01 are shown. Size of the circles is proportional to the number of genes present in the annotation in each category (from 10 to 220). This figure is equivalent to fig. 5 except for the normalisation with category 2 instead of category 1 variants.

**FIG. S13**

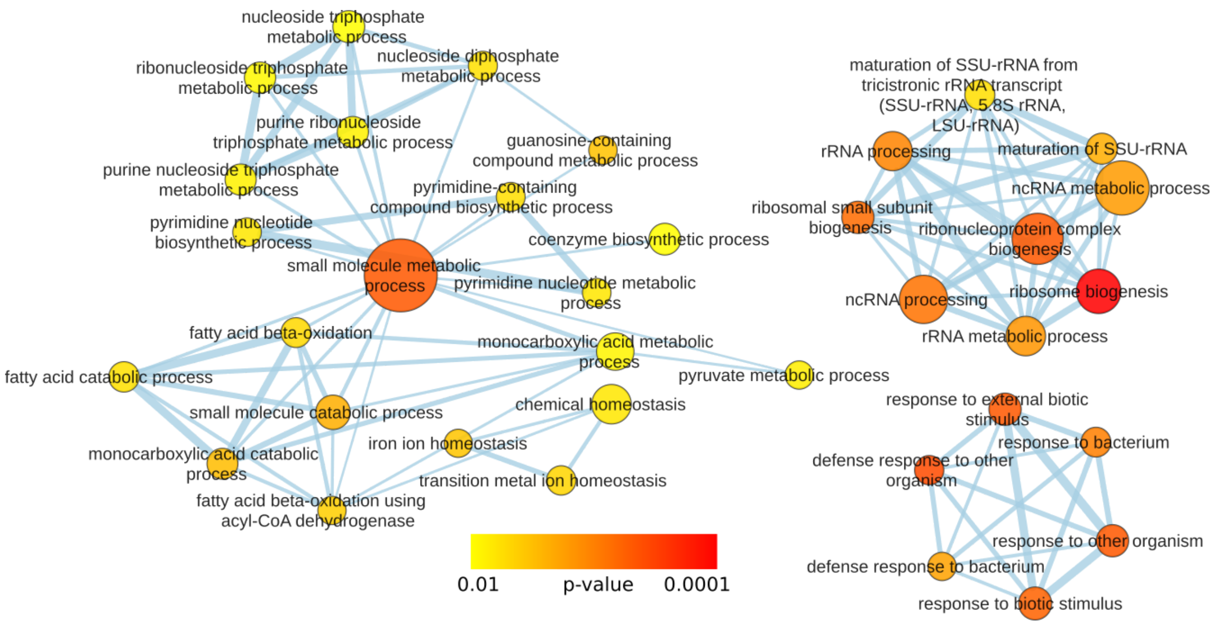


Enrichment of hybridisation-derived variants in biological process gene ontologies (GOs). Genes in the *Oreochromis niloticus* gene annotation were linked to zebrafish GO categories and enrichment of *Malawi* variants differentially fixed between *Victoria* and *Ruaha Blue* (category 3 in table 1) relative to accessible genome length in exonic regions was tested (see Materials and Methods). Categories with enrichment p-values <0.01 are shown. Size of the circles is proportional to the number of genes present in the annotation in each category (from 10 to 220). This figure is equivalent to fig. 5 except for the normalisation by accessible genome length instead of category 1 variants.

---

**FIG. S14**

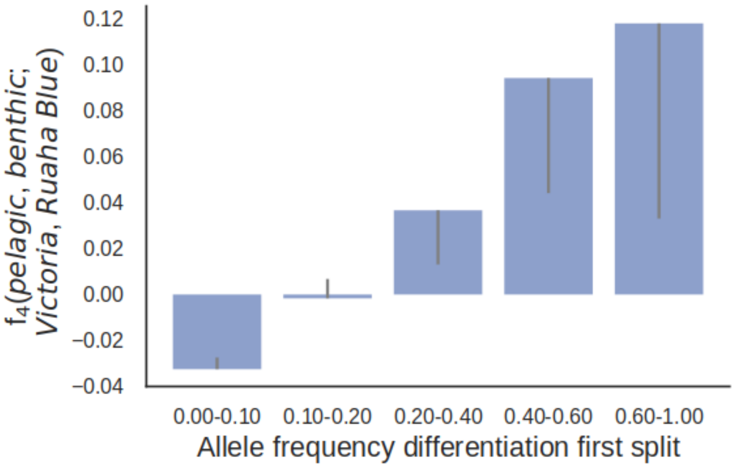


Allele sharing of lineages descending from *Malawi* first split with Victoria/*Ruaha Blue*. $f_4$ admixture ratio measuring the amount of *Victoria* ancestry in *benthic* relative to *pelagic* for hybridisation-derived variants. Error bars correspond to three jackknifing standard deviations and are only drawn towards the origin. Error bars that do not overlap the origin indicate to block-jackknifing significance p<0.01 (all bins except 0.10-0.20).

**FIG. S15**

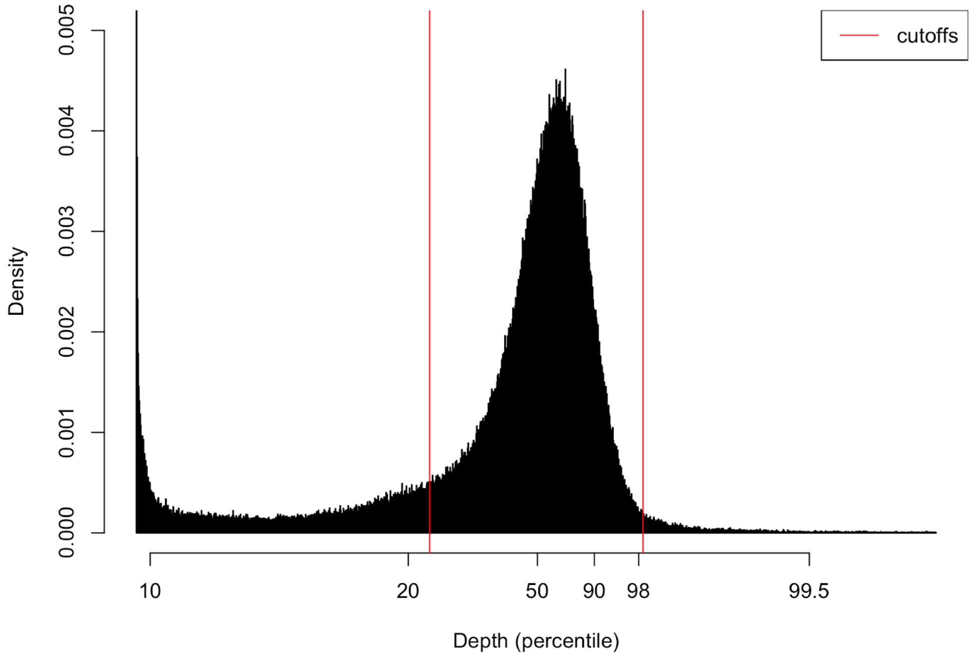


The cutoffs for filtering on overall depth (DP annotation in the VCF file) were selected by manually examining the distribution.

### Supplementary Tables

Supplementary table 1 is available as separate .xlsx document.
