## Supplementary table 1 for "Ancestral hybridisation facilitated species diversification in the Lake Malawi cichlid fish adaptive radiation"

| Image | Image notes and credit | NCBI taxon | Publication name | Reference | Sample collection date | Location | Latitude | Longitude | Biosample | Bioproject | Sample name in VCF file | Sequencing coverage |
|---|---|---|---|---|---|---|---|---|---|---|---|---|
| 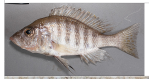 | Image not the actual sample. | Alticorpus geoffreyi | Alticorpus geoffreyi | Malinsky et al. 2018 | | Lake Malawi | | | SAMEA1904331 | PRJEB1254 | Alticorpus_geoffreyi | 12.1 |
| 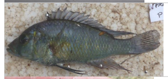 | | Astatotilapia bloyeti | Astatotilapia bloyeti | Malinsky et al. 2018 | 12/08/2015 | Lake Kumba | -4.80628 | 38.62163 | SAMEA4033334 | PRJEB15289 | Astatotilapia_bloyeti_MG_P2C8 | 17 |
| 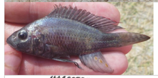 | | Astatotilapia bloyeti | Astatotilapia bloyeti 2 | Malinsky et al. 2018 | 16/08/2015 | Lake Burungi | -3.91563 | 35.86081 | SAMEA4033333 | PRJEB15289 | Astatotilapia_bloyeti_MG_A17_13 | 15.8 |
| 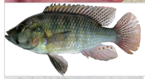 | | Astatotilapia bloyeti | Astatotilapia bloyeti 3 | This paper | | Uvinza, Tanzania (Malagarasi) | -5.109444 | 30.393611 | In submission. Sample and project numbers pending. | In submission. Sample and project numbers pending. | Haplochromis_paludinosus | 7.9 |
| 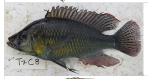 | | Astatotilapia bloyeti | Astatotilapia bloyeti 4 | Malinsky et al. 2018 | 25/7/2015 | Lake Nala, near Kilosa, Wami System | -6.945 | 36.937 | SAMEA4033341 | PRJEB15289 | Astatotilapia_bloyeti_GT_T7C8 | 10 |
| 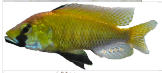 | Image not the actual sample. By Emília Santos. | Astatotilapia calliptera | Astatotilapia calliptera Malawi | Malinsky et al. 2018 | | Lake Malawi | | | SAMEA1920092 | PRJEB1254 | A_calliptera_Salima_Father | 10.4 |
| 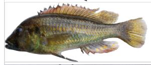 | Image not the actual sample. | Astatotilapia | Astatotilapia calliptera Massoko | Malinsky et al. 2018 | | Lake Massoko | | | SAMEA1877464 | PRJEB1254 | Massoko_benthic_HC_1 | 10.5 |
| 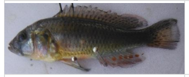 | | Haplochromis tweddlei | Astatotilapia gigliolii | Malinsky et al. 2018 | 29/01/2014 | Ruvu, Mindu Dam | -6.865 | 37.61 | SAMEA4033340 | PRJEB15289 | Astatotilapia_tweddlei_GT_2D10 | 9.8 |
| no image | | Haplochromis tweddlei | Astatotilapia gigliolii 2 | This paper | | Aquarium specimen, possibly from area between Ifakara and Kidatu | | | In submission. Sample and project numbers pending. | In submission. Sample and project numbers pending. | HaplochromisKilossana | 9.9 |
| 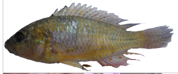 | | Haplochromis tweddlei | Astatotilapia gigliolii 3 | Malinsky et al. 2018 | 29/01/2014 | Ruaha, Kidatu | -7.662 | 36.978 | SAMEA4033339 | PRJEB15289 | Astatotilapia_tweddlei_GT_2I2 | 16 |
| 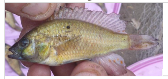 | | Haplochromis tweddlei | Astatotilapia gigliolii 4 | Malinsky et al. 2018 | 19/08/2013 | Kitele Lake | -10.3595 | 39.77558333 | SAMEA4033332 | PRJEB15289 | Astatotilapia_tweddlei_MG174A | 14.6 |
| 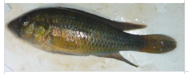 | | Haplochromis tweddlei | Astatotilapia gigliolii 5 | Malinsky et al. 2018 | 09/09/2012 | Near Songea | -10.62533333 | 35.65338889 | SAMEA4033331 | PRJEB15289 | Astatotilapia_tweddlei_MG221 | 11.6 |
| no image | | Haplochromis tweddlei | Astatotilapia gigliolii 6 | Malinsky et al. 2018 | 18/05/2009 | Nkhokwe, Lake Chiuta | -14.725 | 35.78333333 | SAMEA3388867 | PRJEB1254 | Astatotilapia_tweddlei | 14.6 |
| 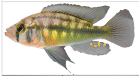 | | Haplochromis sp. Chipwa | Astatotilapia sp. "chipwa" | This paper | | Kalambo River Delta, Chipwa Village, Zambia | -8.6017 | 31.186709 | In submission. Sample and project numbers pending. | In submission. Sample and project numbers pending. | Haplochromis_sp_chipwa | 8.1 |
| 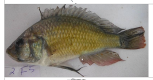 | Photo mislabeled. | New taxon (submitted to NCBI) | Astatotilapia sp. "longfin yellow" | This paper | 29/01/2014 | Kidatu, Gt Ruaha, Rufiji System | -7.662 | 36.978 | In submission. Sample and project numbers pending. | In submission. Sample and project numbers pending. | Astatotilapia_ruaha_yellow_GT_2I5 | 14.7 |
| 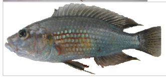 | | Astatotilapia sp. 'Ruaha | Astatotilapia sp. "ruaha blue" 2 | This paper | 29/01/2014 | Kidatu, Gt Ruaha, Rufiji System | -7.662 | 36.978 | In submission. Sample and project numbers pending. | In submission. Sample and project numbers pending. | Astatotilapia_ruaha_blue_GT_2J2 | 13.6 |

| Image | Image notes and credit | NCBI taxon | Publication name | Reference | Sample collection date | Location | Latitude | Longitude | Biosample | Bioproject | Sample name in VCF file | Sequencing coverage |
|---|---|---|---|---|---|---|---|---|---|---|---|---|
| 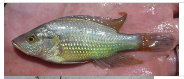 | | Astatotilapia sp. 'Ruaha | Astatotilapia sp. "ruaha blue" | Malinsky et al. 2018 | 05/09/2012 | Rujewa, fish ponds | -8.70825 | 34.38822222 | SAMEA2661249 | PRJEB1254 | Astatotilapia_rujewa | 19.2 |
| 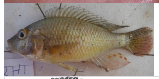 | | New taxon (submitted to NCBI) | Astatotilapia sp. "ruaha red cheek" | This paper | 29/01/2014 | Kidatu, Gt Ruaha, Rufiji System | -7.662 | 36.978 | In submission. Sample and project numbers pending. | In submission. Sample and project numbers pending. | Astatotilapia_ruaha_yellow_GT_2H1 | 10.7 |
| 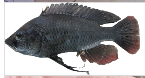 | | Astatotilapia sp. 'Blue Rufiji' | Astatotilapia sp. "rufiji blue" | This paper | 05/06/2015 | Lake Mansi | -7.276 | 39.567 | In submission. Sample and project numbers pending. | In submission. Sample and project numbers pending. | Astatotilapia_rufiji_blue_MG_T5B3 | 14 |
| 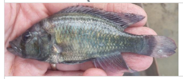 | | Astatotilapia sp. 'Blue Rufiji' | Astatotilapia sp. "rufiji blue" 2 | This paper | 20/08/2013 | Oxbow Lake, Utete | -7.990861111 | 38.74925 | In submission. Sample and project numbers pending. | In submission. Sample and project numbers pending. | Astatotilapia_rufiji_blue_MG_188A | 14 |
| 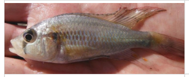 | | New taxon (submitted to NCBI) | Astatotilapia sp. "rukwa blue" | This paper | 03/09/2012 | Lake Rukwa | -8.397472222 | 32.90183333 | In submission. Sample and project numbers pending. | In submission. Sample and project numbers pending. | Astatotilapia_rukwa_blue_MG_49A | 15.7 |
| 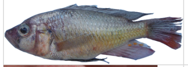 | | New taxon (submitted to NCBI) | Astatotilapia sp. "rukwa large yellow" | This paper | 02/09/2012 | Mkwajundi Market, Lake Rukwa | -8.4431 | 33.013232 | In submission. Sample and project numbers pending. | In submission. Sample and project numbers pending. | Astatotilapia_rukwa_large_yellow_MG_203 | 14.6 |
| 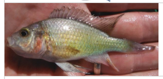 | | New taxon (submitted to NCBI) | Astatotilapia sp. "rukwa sunset" | This paper | 03/09/2012 | Lake Rukwa | -8.397472222 | 32.90183333 | In submission. Sample and project numbers pending. | In submission. Sample and project numbers pending. | Astatotilapia_rukwa_sunset_MG_57A | 13.1 |
| no image | | New taxon (submitted to NCBI) | Copadichromis likomae | Malinsky et al. 2018 | | Lake Malawi | | | In submission. Sample and project numbers pending. | In submission. Sample and project numbers pending. | Copadichromis_likomae | 16.1 |
| 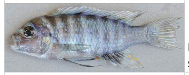 | Image not the actual sample. | Cynotilapia afra | Cynotilapia afra | Malinsky et al. 2018 | | Lake Malawi | | | SAMEA2661253 | PRJEB1254 | Cynotilapia_afra | 15.5 |
| 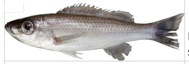 | Image not the actual sample. | Diplotaxodon limnothrissa | Diplotaxodon limnothrissa | Malinsky et al. 2018 | | Lake Malawi | | | SAMEA2661260 | PRJEB1254 | Diplotaxodon_limnothrissa | 16.2 |
| no image | | New taxon (submitted to NCBI) | Diplotaxodon sp. "macrops ngulube" | Malinsky et al. 2018 | | Lake Malawi | | | In submission. Sample and project numbers pending. | In submission. Sample and project numbers pending. | Diplotaxodon_ngulube | 14.6 |
| 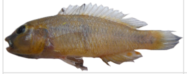 | | Haplochromis vanheusdeni | Haplochromis vanheusdeni | This paper | 30/01/2014 | Kidatu, Gt Ruaha, Rufiji System | -7.662 | 36.978 | In submission. Sample and project numbers pending. | In submission. Sample and project numbers pending. | Haplochromis_vanheusdeni_GT_3A1 | 14.3 |
| no image | | Haplochromis vanheusdeni | Haplochromis vanheusdeni 2 | This paper | | Ruaha River | -7.808222 | 36.896556 | In submission. Sample and project numbers pending. | In submission. Sample and project numbers pending. | Haplochromis_vanheusdeni | 8.7 |
| 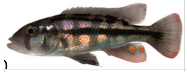 | McGee et al. 2016 | Haplochromis sp. 'checkmate' | Harpagochromis sp. "checkmate" | McGee et al. 2016 | | Lake Victoria | | | SAMN04158999 | PRJNA298405 | Harpagochromis_checkmate_Victoria | 10 |
| 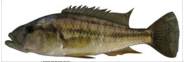 | McGee et al. 2016 | Haplochromis vittatus | Harpagochromis vittatus | McGee et al. 2016 | | Lake Kivu | | | SAMN04158996 | PRJNA298405 | Harpagochromis_vittatus_Kivu | 16.3 |
| 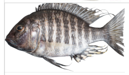 | Image not the actual sample. | Lethrinops gossei | Lethrinops gossei | Malinsky et al. 2018 | | Lake Malawi | | | SAMEA1877484 | PRJEB1254 | Lethrinops_gossei | 12.2 |

| Image | Image notes and credit | NCBI taxon | Publication name | Reference | Sample collection date | Location | Latitude | Longitude | Biosample | Bioproject | Sample name in VCF file | Sequencing coverage |
|---|---|---|---|---|---|---|---|---|---|---|---|---|
| 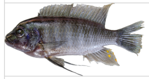 | Image not the actual sample. | Maylandia zebra | Maylandia zebra | Malinsky et al. 2018 | | Lake Malawi | | | SAMEA3388874 | PRJEB1254 | Metriaclima_zebra | 14.3 |
| 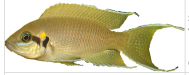 | Image not the actual sample. Modified from Wikipedia. By Przemysław Malkowski. CC BY-SA 3.0 | Neolamprologus brichardi | Neolamprologus brichardi | Brawand et al. 2014 | | Lake Tanganyika | | | SAMN00139653 | PRJNA60365 | N_brichardi2 | 14.9 |
| no image | | Otopharynx lithobates | Otopharynx lithobates | Malinsky et al. 2018 | | Lake Malawi | | | SAMEA3388872 | PRJEB1254 | Otopharynx_lithobates | 13.8 |
| 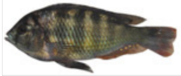 | McGee et al. 2016 | Haplochromis flavus | Paralabidochromis flavus | McGee et al. 2016 | | Lake Victoria | | | SAMN04158998 | PRJNA298405 | Paralabidochromis_flavus_Victoria | 15.3 |
| 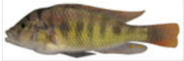 | McGee et al. 2016 | Haplochromis paucidens | Paralabidochromis paucidens | McGee et al. 2016 | | Lake Kivu | | | SAMN04158997 | PRJNA298405 | Paralabidochromis_paucidens_Kivu | 16.1 |
| 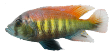 | Image not the actual sample. Modified from Wikipedia. By Oliver Selz. CC BY-SA 3.0 | Pundamilia nyererei | Pundamilia nyererei | Brawand et al. 2014 | | Lake Victoria | | | SAMN00788761 | PRJNA83153 | P_nyererei2 | 17.4 |
| 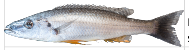 | Image not the actual sample. | Rhamphochromis longiceps | Rhamphochromis longiceps | Malinsky et al. 2018 | | Lake Malawi | | | SAMEA3388853 | PRJEB1254 | Rhamphochromis_longiceps | 14.9 |
| 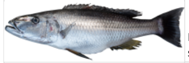 | Image not the actual sample. | Rhamphochromis woodi | Rhamphochromis woodi | Malinsky et al. 2018 | | Lake Malawi | | | SAMEA3388856 | PRJEB1254 | Rhamphochromis_woodi | 13.6 |
| 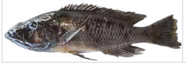 | Image not the actual sample. | Tyrannochromis nigriventer | Tyrannochromis nigriventer | Malinsky et al. 2018 | | Lake Malawi | | | SAMEA3388855 | PRJEB1254 | Tyrannochromis_nigriventer | 14.4 |
